# Super-enhancer-driven *CACNA2D2* is an EWSR1::WT1 signature gene encoding a diagnostic marker for desmoplastic small round cell tumor (DSRCT)

**DOI:** 10.1101/2024.07.17.603708

**Authors:** Florian H. Geyer, Alina Ritter, Seneca Kinn-Gurzo, Tobias Faehling, Jing Li, Armin Jarosch, Carine Ngo, Endrit Vinca, Karim Aljakouch, Azhar Orynbek, Shunya Ohmura, Thomas Kirchner, Roland Imle, Laura Romero-Pérez, Stefanie Bertram, Enrique de Álava, Sophie Postel-Vilnay, Ana Banito, Martin Sill, Yvonne M.H. Versleijen-Jonkers, Benjamin F.B. Mayer, Martin Ebinger, Monika Sparber-Sauer, Sabine Stegmaier, Daniel Baumhoer, Wolfgang Hartmann, Jeroen Krijgsveld, David Horst, Olivier Delattre, Patrick J. Grohar, Thomas G. P. Grünewald, Florencia Cidre-Aranaz

## Abstract

Desmoplastic small round cell tumor (DSRCT) is a highly aggressive cancer predominantly occurring in male adolescents and young adults. The lack of a comprehensive understanding on the biology of the disease is paralleled by its dismal survival rates (5–20%). To overcome this challenge, we first identified and prioritized urgently needed resources for clinicians and researchers. Thus, we established genome-wide single-cell RNA-sequencing and bulk proteomic data of in vitro and in vivo-generated knockdown models of the pathognomonic DSRCT fusion oncoprotein (EWSR1::WT1) and combined them with an original systems-biology-based pipeline including patient data and the largest histology collection of DSRCTs and morphological mimics available to date. These novel tools were enriched with curated public datasets including patient- and cell line-derived ChIP-seq, bulk and single-cell RNA-seq studies resulting in a multi-model and multi-omic toolbox for discovery analyses. As a proof of concept, our approach revealed the alpha-2/delta subunit of the voltage-dependent calcium channel complex, CACNA2D2, as a highly overexpressed, super-enhancer driven, direct target of EWSR1::WT1. Single-cell and bulk-level analyses of patient samples and xenografted cell lines highlighted CACNA2D2 as a critical component of our newly established EWSR1::WT1 oncogenic signature, that can be employed to robustly identify DSRCT in reference sets. Finally, we show that CACNA2D2 is a highly sensitive and specific single biomarker for fast, simple, and cost-efficient diagnosis of DSRCT. Collectively, we establish a large-scale multi-omics dataset for this devastating disease and provide a blueprint of how such toolbox can be used to identify new and clinically relevant diagnostic markers, which may significantly reduce misdiagnoses, and thus improve patient care.

## INTRODUCTION

Desmoplastic small round cell tumor (DSRCT) is an ultra-rare and highly aggressive cancer entity^1,2^, which typically develops at sites lined by mesothelium, such as the abdominal cavity of mostly male adolescents and young adults (male to female ratio: 5:1)^1^. The 5-year survival rate of only 5–20% can be attributed to a lack of effective therapeutic options and standard treatment protocols^1^.

DSRCT is of unknown origin^3^ and genetically defined by a chromosomal translocation fusing the N-terminus of EWS RNA binding protein 1 (*EWSR1*) to the C-terminus of Wilms tumor protein (*WT1)*^4^. The encoded pathognomonic fusion protein, EWSR1::WT1, is a potent transcription factor and the only known driver of oncogenic transformation in DSRCT^5^.

Despite the recognition of DSRCT as a distinct entity for several decades^6^, large-scale studies have remained scarce due to the ultra-rare nature of the disease. Though in recent years several efforts have been made to advance our understanding of DSRCT^7–12,5,13–16^, the resulting datasets are often fragmented and insufficient for a comprehensive characterization of the complexity of this disease. For instance, to date the few available gene expression datasets exploring DSRCT have been derived from in vitro (often individually) grown cell lines and exclusively at (predominantly bulk) epigenetic or transcriptomic levels, including ChIP-seq, Hi-ChIP, and RNA-seq/microarray-based profiling^5,7,8,14,17,18^. All in all, there is a lack of a) available data from systematically multi-omic-profiled and physiologically-grown subcutaneously or orthotopically xenografted cell lines, especially at proteomic and single cell levels, that could better recapitulate the particularities of this disease; and b) integration of already existing public datasets into a comprehensive resource that can be further exploited for future research on DSRCT.

Similarly, reliable and accurate diagnosis of DSRCT using immunohistochemistry (IHC) has remained a major clinical challenge because of its heterogenous and often indistinctive morphology^2,19,6^. In addition, due to the rarity of the disease, only small collections of histological DSRCT patient samples have remained geographically scattered in different institutions, hindering the exploration and validation of new markers in a larger cohort more widely representative of the spectrum of DSRCT patients.

In this study, we generated novel multi-omic datasets – including proteomic and genome-wide, single-cell RNA sequencing of in vivo xenografted DSRCT cell lines with conditional knockdowns of EWSR1::WT1 – to allow a deeper understanding of the disease and provide the scientific community with a large, valuable resource for future research. In addition, we enriched these data with curated public datasets including patient- and cell line-derived ChIP-seq, bulk and single-cell RNA-seq studies, together with the largest DSRCT histological cohort to date amenable to biomarker validation, resulting in a multi-model and multi-omic toolbox for discovery analyses. By harnessing this comprehensive toolbox, we provided a blueprint to identify new and clinically relevant markers, such as the super-enhancer (SE)-driven EWSR1::WT1 core signature gene *CACNA2D2*, whose expression can be used as a highly specific and sensitive single IHC-marker for robust diagnosis of DSRCT.

## METHODS

### Provenience of cell lines and cell culture conditions

Human cell lines were provided by the following repositories and/or sources: MeT-5A human mesothelial cells (ATTC number: CRL-9444) and HEK293T (ATTC number: CRL-3216) cells were purchased from the American Type Culture Collection (ATCC, Manassas, VA, USA). Human DSRCT cell line JN-DSRCT-1 was provided by Dr. Mikiko Aoki (Fukuoka University, Fukuoka, Japan), SK-DSRCT2 was kindly provided by Dr. Marc Ladanyi (Memorial Sloan Kettering Cancer Center, New York, NY, USA), and BER cells were obtained from the Christus Stehlin Foundation for Cancer Research^20^. All cell lines were cultured in RPMI-1640 medium (Sigma-Aldrich, St Louis, MO, USA) supplemented with 10% tetracycline-free fetal calf serum (FCS; Sigma-Aldrich) and 10,000 U/ml penicillin and 10 mg/ml streptomycin (Sigma-Aldrich) at 37°C with 5% CO2 in a humidified atmosphere. Cell lines were routinely tested for the absence of mycoplasma contamination using nested PCR. Additionally, cell line identity was regularly verified through Short Tandem Repeat (STR) profiling or Single Nucleotide Polymorphism (SNP) profiling.

### Generation of Doxycycline-inducible short hairpin RNA (shRNA) expressing cells

Human DSRCT cell lines JN-DSRCT-1 and SK-DSRCT2 were transduced with lentiviruses encoding the TET-pLKO-puro vector (Plasmid #21915, Addgene, Watertown, MA, USA) containing a puromycin resistance and doxycycline (DOX)-inducible expression cassette of short hairpin RNAs (shRNAs) against *WT1* (shWT1) or a non-targeting control shRNA (shCtrl)^21^. DOX-inducible vectors were generated as described before^22^ using a non-targeting shRNA (shCtrl) or two shRNAs against *WT1* as inserts, targeting either the coding sequence (shWT1-CDS) or the 3’-UTR (shWT1-UTR) of the *WT1* RNA transcript (**Suppl. Table 1**). Vectors were amplified in NEB^®^ Stable Competent *E. coli* (New England Biolabs, Ipswich, MA, USA) under antibiotic selection with Ampicillin (Sigma-Aldrich) at 100 μg/ml in LB Broth (Carl Roth GmbH + Co. KG, Karlsruhe, Germany) at 30°C and 180 rpm. Integrated shRNA was verified by Sanger-sequencing (sequencing primer sequence: 5’-GGCAGGGATATTCACCATTAT-3’). Lentiviral particles were generated in HEK293T cells using psPAX2 (Plasmid #12260, Addgene) and pMD2.G (Plasmid #12259, Addgene) packaging plasmids. Virus-containing supernatant was collected to infect the human DSRCT cell lines. Infected cells were selected with 1 µg/ml puromycin (Sigma-Aldrich). The shRNA expression for *WT1* knockdown in DSRCT cells was achieved by adding 1 µg/ml DOX (Sigma-Aldrich) every 72 h to the cell culture medium. Generated cell lines were designated JN-DSRCT-1/TR/shCtrl, JN-DSRCT-1/TR/shWT1-CDS, JN-DSRCT-1/TR/shWT1-UTR, SK-DSRCT2/TR/shCtrl, SK-DSRCT2/TR/shWT1-CDS, and SK-DSRCT2/TR/shWT1-UTR.

### Transient transfection using small interfering RNAs (siRNA)

Two siRNA sequences, each targeting one side of the EWSR1::WT1 fusion breakpoint (**Suppl. Table 1**), were used to selectively suppress EWSR1::WT1 without impacting wildtype EWSR1 or WT1 expression as previously described^5^. Pooled siRNAs or non-targeting siRNA were complexed with 3 μl Lipofectamine RNAiMax Transfection Reagent (Thermo Scientific, Dreieich, Germany) for 30 min at 37 °C temperature. siRNAs were added to 5×10^5^ BER DSRCT cells with cell culture medium at a final concentration of 2 nmol/l. After incubation for 48h at 37°C, cells were washed with PBS (Thermo Scientific) and either used for either RNA or protein extraction.

### CRISPR-Cas9 knock-in establishment and PROTAC-mediated degradation

Clustered regulatory interspaced short palindromic repeats (CRISPR)-Cas9 system was used to knock-in the HaloTag-HibiT-Tag at the endogenous locus of *EWSR1::WT1*^23,24^. First, custom crRNA targeting the C-terminus of *EWSR1::WT1* (Integrated DNA Technologies, Inc., Coralville, IA, USA) was annealed with tracrRNA (Integrated DNA Technologies). Next, Alt-R™ S.p. HiFi Cas9 Nuclease enzyme (Integrated DNA Technologies) was added to the annealed sgRNA to form a ribonucleoprotein (RNP) complex. RNP complex was delivered into 1×10^6^ SK-DSRCT2 cells via transient transfection using the SF Cell Line 4D-Nucleofector™ X Kit and the 4D-NucleofectorTM X Unit (Lonza, Basel, Switzerland) using the pulse code DS-1337. Isogenic cell clones were then retrieved by serial dilution and knock-in was validated via PCR and subsequent Sanger-sequencing. The generated cell line was designated SK-DSRCT2-endo-WT1-HaloTag. crRNA and PCR primer sequences are provided in **Suppl. Table 1**. To induce the post-translational knock down of tagged EWSR1::WT1, a PROTAC (Promega Corporation, Madison, WI, USA) targeting the HaloTag was added to cell culture medium at 1 μmol/l for 24 h. Knock down efficiency was then tested with Nano-Glo^®^ HiBiT Lytic Detection System (Promega), or western blot.

### Generation of DOX-inducible EWSR1::WT1 expressing mesothelial cells

Human mesothelial cell line MeT-5A was transduced with lentiviruses encoding pTPin overexpression vector containing a puromycin resistance and DOX-inducible expression cassette of *EWSR1::WT1^−KTS^* as described before^25^. The DOX-inducible vector was generated by enzymatic digestion of the vector with AgeI (New England Biolabs) and NotI (New England Biolabs) restriction enzymes. Total RNA from SK-DSRCT2 wild-type cell line was reverse transcribed using SuperScript™ IV Reverse Transcriptase (Thermo Scientific) and used to amplify EWSR1::WT1 mRNA sequence. EWSR1::WT1 cDNA sequence was amplified with primers adding overhangs for AgeI and NotI restriction enzymes. PCR product was digested and ligated into the digested pTPin vector. PCR primer sequences are provided in **Suppl. Table 1.** Vectors were amplified in NEB^®^ Stable Competent *E. coli* (New England Biolabs) under antibiotic selection with Ampicillin (Sigma-Aldrich) at 100 μg/ml in LB Broth (Carl Roth) at 30°C and 180 rpm for 18 h. The integrated transgene was verified by Sanger-sequencing. Lentiviral particles were generated in HEK293T cells using psPAX2 (Plasmid #12260, Addgene) and pMD2.G (Plasmid #12259, Addgene) packaging plasmids. Virus-containing supernatant was collected to infect the mesothelial cell line. Infected cells were selected with 1 µg/ml puromycin dihydrochloride (Sigma-Aldrich). Expression of *EWSR1::WT1* was achieved by adding 1 µg/ml DOX (Sigma-Aldrich) every 72h to the cell culture medium. The generated cell line was designated MeT-5A-EWSR1::WT1^−KTS^.

### RNA extraction, reverse transcription, and quantitative real-time PCR (qRT-PCR)

Total RNA was isolated using the NucleoSpin RNA kit (MACHEREY-NAGEL GmbH & Co. KG, Düren, Germany). 1 µg of total RNA was reverse-transcribed using High-Capacity cDNA Reverse Transcription Kit (Applied Biosystems, Waltham, MA, USA). qRT-PCR reactions (final volume 15 µl) were performed using SYBR™ Select Master Mix for CFX (Applied Biosystems, Waltham, MA, USA) mixed with diluted cDNA (1:10) and 0.5 µM forward and reverse primer on a BioRad CFX Connect instrument and analyzed using CFX Maestro 2.0 software (Bio-Rad Laboratories GmbH, Feldkirchen, Germany). Gene expression values were calculated using the ^ΔΔ^Ct method relative to the housekeeping gene *RPLP0* as an internal control. Oligonucleotides were purchased from Merck KGaA (Darmstadt, Germany) (**Suppl. Table 1**). Thermal conditions for qRT-PCR were as follows: UDG-Activation at 50°C for 2 min, heat activation at 95°C for 2 min, DNA denaturation at 95°C for 15 s, annealing and elongation at 60°C for 60 s (40 cycles).

### Analysis of published chromatin immuno-precipitation followed by high-throughput DNA sequencing (ChIP-seq) data and binding motif analysis

Preprocessed ChIP-seq data from JN-DSRCT-1 cell line was downloaded from the Gene Expression Omnibus data repository with accession codes GSE156277 and GSE212977. ChIP-seq data from four primary DSRCT specimens and MeT-5A cell line were downloaded from GEO (accession code: GSE212977). Data was displayed in the UCSC genome browser as custom tracks. The samples analyzed in this study are listed in **Suppl. Table 2**. EWSR1::WT1 binding motifs were identified by analyzing previously published genomic positions of EWSR1::WT1 binding sites in Browser Extensible Data (BED) format^14^ with HOMER’s findMotifsGenome.pl wrapper performing the HOMER motif discovery algorithm with hg19 reference genome^26,27^. Find Individual Motif Occurrences (FIMO) analysis was performed as part of the MEME Suite^28^. FIMO was fed with genomic DNA sequences of interest extracted from the UCSC genome browser (hg19) in FASTA format, and the top five de novo and known binding motifs detected in ChIP-seq data of EWSR1::WT1 binding in JN-DSRCT-1 cell line. FIMO analysis results were downloaded in General Feature Format (version 3) and uploaded as a custom track to the UCSC genome browser.

### Analysis of published Hi-ChIP data

Hi-ChIP datasets used in this study were downloaded from the Gene Expression Omnibus (GEO) repository (accession code: GSE212978). Published preprocessed data was filtered for high-quality reads (read quality>30). The filtered high-quality reads were plotted in an x-y scatterplot to visualize the spatial interactions between genomic loci captured by the Hi-ChIP experiment, where the x and y coordinates represent the genomic positions of the interacting loci, and the density of points reflects the frequency of interactions. Highlighted areas include *CACNA2D2* enhancer region in hg19 reference genome (chr3:50,514,500–50,524,800) and *CACNA2D2* promoter region (chr3:50,538,700–50,542,200).

### Super-enhancer analysis

Raw ChIP-seq data was downloaded as *.fastq files from the Sequence Read Archive (SRA)^29^ and aligned to the hg19 reference genome using BWA-MEM algorithm^30^. SRA run IDs are provided in **Suppl. Table 3**. Resulting *.sam files were converted to *.bam files, sorted and indexed using samtools^31^. H3K27ac *.bam files and corresponding DNA input *.bam files were used to call significant ChIP-seq peaks with macs2 algorithm^32^. Resulting *.bed file and corresponding sorted and indexed *.bam files were used to identify SE using Rank ordering of SE (ROSE) algorithm^33,34^. Identified enhancers were filtered for known regions on the reference genome and sorted according to their H3K27ac density values.

### Tumor dissociation for single-cell RNA-sequencing (scRNA-seq)

Orthotopically xenografted tumors from SK-DSRCT2 and JN-DSRCT-1 cell lines with shRNA-knockdown of the EWSR1::WT1 were processed into single-cell suspensions as previously described^35^. Briefly, tumor sections (≤ 200 mg) were harvested, rapidly cooled in ice-cold PBS (Thermo Scientific), and kept on ice until further processing. Sections were minced with a scalpel and transferred to gentleMACS C tubes (Miltenyi Biotec, Bergisch-Gladbach, Germany) with 5 mL of ice-cold protease cocktail, consisting of cold-active Bacillus licheniformis protease (10 mg/mL, Sigma-Aldrich), calcium chloride (5 mM, Sigma-Aldrich), DNase I (125 U/mL, Roche, Basel, Switzerland) in PBS (Thermo Scientific) and incubated for 10 min at 4°C with rocking. Then, the C tubes were placed in a gentleMACS Dissociator (Miltenyi Biotec, Bergisch-Gladbach, Germany) at 4°C and the m_brain_03 program was run twice. After incubation at 4°C for 5 min, the suspension was filtered through a 70-µm cell strainer (Greiner Bio-One, Kremsmünster, Austria) and the protease was stopped by addition of 10 mL ice-cold Hanks’ Balanced Salt Solution (HBSS, Corning, Corning, NY, USA) supplemented with 10% tetracycline-free fetal calf serum (Sigma-Aldrich). After centrifugation at 300 × g for 5 min at 4°C, the supernatant was discarded, and the pelleted cells were resuspended in ice-cold HBSS supplemented with 2% bovine serum albumin (PAN-Biotech GmbH, Aidenbach, Germany). Cells were first incubated with Human TruStain FcX™ (Fc Receptor Blocking Solution) (BioLegend, California, CA, USA) and TruStain FcX™ PLUS (anti-mouse CD16/32) antibody (BioLegend) on ice for 5 min. Subsequently, APC anti-human CD36 (BioLegend), Brilliant Violet 711™ anti-human CD63 (BioLegend), and FITC anti-mouse CD90.2 (Thy1.2) (BioLegend) antibodies were added at a 1:20 dilution and incubated for 15 min on ice. Cells positive for CD36 and CD63 and negative for CD90.2 were defined as DSRCT and 96 cells per tumor sample were sorted into a DNA low-binding 384-well plate containing 1.2 µL of lysis mix per well (0.095% Triton X-100 (Sigma-Aldrich), 1 U/µL Recombinant RNase Inhibitor (Takara Bio, San Jose, CA, USA), 2,5 mM dNTPs (Thermo Fisher), 2.5 µM Oligo-dT primer (5’-AAGCAGTGGTATCAACGCAGAGTACTTTTTTTTTTTTTTTTTTTTTTTTTTTT-TTVN-3’, Sigma-Aldrich)) using FACSAria Fusion cytometer (BD Biosciences, Franklin Lakes, NJ, USA). scRNA-seq using a modified SMART-seq2 method (V2.5)^36^. Liquid handling was assisted by the Mantis (Formulatrix, Bedford, MA, USA), BRAVO (Agilent, Santa Clara, CA, USA) and Mosquito LV (SPT Labtech, Melbourn, UK) liquid handling platforms. Lysed cells were thawed and incubated for 3 min at 72°C to complete lysis. After 2 min incubation on ice, reverse transcription and template switching mix (10 U/µL Maxima H-reverse transcriptase (Thermo Fisher), 1.4 U/µL Recombinant RNase Inhibitor (Takara Bio), 1X Maxima H-buffer (Thermo Fisher), 4,5 µM template switching oligo (AAGCAGTGGTATCAACGCAGAGTACATrGrG+G, Eurogenetec and 11,25% poly-ethylenglycol 8000 (Sigma-Aldrich) in nuclease-free water (Thermo Fisher)) were added to each well and the plates were incubated for 90 min at 42°C, followed by an enzyme inactivation at 70°C for 15 min. PCR amplification with 21 cycles was conducted by adding 3 µL KAPA HiFi HotStart ReadyMix (Roche) and 0.05 µL IS PCR primer (10 µM) to each well and incubating according to the manufacturer’s protocol with 67°C annealing temperature and 6 min extension. After clean-up with 5 µL AmpureXP beads (Beckman-Coulter, Brea, CA, USA) at a beads ratio of 0.8, quality control was conducted with Tapestation D5000 reagents (Agilent) and cDNA was diluted to 1–3 ng/µL. Fragmentation of full-length cDNA was performed using 1.2 µL Tn5 tagmentase in tagmentation buffer per well (10 mM Tris-HCl pH 7.5 (Serva electrophoresis, Heidelberg, Germany), 10 mM magnesium chloride (Thermo Fisher), 50% dimethylformamide (Sigma-Aldrich)) with 0.4 µL cDNA by incubating for 10 min at 55°C. After stopping the incubation by adding 0.4 µL sodium dodecyl sulfate (Sigma-Aldrich), barcoding oligos with Illumina sequencing barcodes were added, and library amplification was conducted by adding 2.7 µL KAPA HiFi HotStart ReadyMix (Roche) with 0.3 µL dimethyl-sulfoxide (Sigma-Aldrich) and incubating according to the manufacturer’s protocol with 12 cycles of 63°C annealing temperature and 30 s extension time. After pooling and clean-up with AmpureXP beads (Beckman-Coulter) at a beads ratio of 0.9 according to manufacturer’s instructions, quality control was conducted with Tapestation D1000 reagents (Agilent) and the pooled library was diluted to 10 nM for sequencing on the NextSeq 550 sequencing platform (High output, 75bp single-end, Illumina, San Diego, CA, USA).

### Data analysis of in vivo samples scRNA-seq samples

Quality control of the raw reads was conducted using fastqc, and adapter trimming was done by the Next Generation Sequencing Unit of DKFZ. Alignment was done with the STARsolo method of the Spliced Transcripts Alignments to a Reference tool (STAR 2.7.11b, Alexander Dobin). Quality control, analysis and plotting was done using the R packages Seurat version 5.1.0^37–41^ and SingleCellExperiment version 1.24.0^42^. Quality control was done following the recommendations of R package SeuratWrappers^43^ version 0.3.5 RunMiQC function with a posterior cutoff of 0.8 and the R package scuttle^44^ version 1.12.0 quickPerCellQC function, and by discarding all genes expressed in less than 2 cells. Normalization, scaling and centering was performed using the SCTransform function, considering the potential confounders of ribosomal and mitochondrial gene expression, cell cycle state, expression of housekeeping and dissociation-associated genes^45–48^ as well as the cell line and shRNA sequence. scEWSR1::WT1 and scCACNA2D2 signatures were computed as follows: For scEWSR1::WT1, the FindMarkers function comparing DOX-treated with non-treated cells was used. For scCACNA2D2, the top 100 genes with the highest correlation coefficient with the CACNA2D2 expression values across all cell lines were selected. The signatures were scored using the ssGSEA implementation of the irGSEA R package version 3.2.5^49^.

### Data analysis of scRNA-seq patient samples

The pre-processed and annotated scRNA-seq data from patient samples from Henon et al.^15^ were scored using the ssGSEA implementation of the irGSEA R package and plotted using Seurat functions. All GEO accession codes for the scRNA-seq data analyzed in this study are provided in **Suppl. Table 4.**

### Gene expression microarray normalization

Publicly available and well-curated gene expression data of 654 samples comprising 20 cancer entities and 929 normal tissue samples from 71 normal tissue types generated on GeneChip™ Human Genome U133 Plus 2.0 Array microarrays (Applied Biosystems, Waltham, MA, USA)^50^ were combined with 32 DSRCT samples from our own cohort^51^. GEO accession codes for each dataset analyzed are listed in **Suppl. Table 5**. All microarray *.CEL files were preprocessed (normalized) simultaneously in R version 4.3.0 using the affy package version 1.78.2^52^. For this, the Robust Multi-chip Average (RMA) algorithm, including background adjustment, quantile normalization and summarization was applied^53^. For RMA normalization custom brainarray Chip Description Files (CDF; ENTREZG, v25) were used, yielding one optimized probe set per gene^54^.

### Differential gene/protein expression analyses

Differential gene/protein expression analysis (DEG/DEP analysis) of microarray expression data or protein expression was performed using R limma package version 3.56.2^55^. Preprocessed and normalized expression data was first log2 transformed. Then, DEG/DEP was calculated by applying gene/protein-wise linear modeling fitting, empirical bayes moderation and statistical testing with limma. The resulting *P*-values were adjusted for multiple testing based on false discovery rate (FDR) correction. DEG analysis of RNA-seq data was performed using the standard workflow of R DeSeq2 package version 1.40.2^56^. For this, raw gene counts were retrieved from GEO (GEO accession codes GSE137561, GSE212976 and GSE252051) and lowly expressed genes were removed based on a count threshold of<10 across all samples analyzed. Then read counts were normalized using the DESeq2 method. Differential gene expression testing was done using the Wald test. *P*-values were adjusted for multiple testing using the Benjamini-Hochberg method.

### Fast sample gene-set enrichment analyses (fGSEA) and single sample gene-set enrichment analyses (ssGSEA) and visualization

To identify enriched gene sets within genes that are coexpressed with *CACNA2D2*, genes were ranked according to their Pearson correlation coefficient, and a fast preranked gene-set enrichment analysis (fGSEA) was performed with 10,000 permutations in R using fgsea package version 1.26.0^57^. fGSEA was performed on the hallmark gene set (H) and curated gene sets including canonical pathways from Reactome (C2:CP:REACTOME) from the Human Molecular Signatures Database (MSigDB, version v2023.2.Hs)^58–61^. To visualize fGSEA results, enriched gene sets were filtered for significance (adjusted *P*<0.01; |Normalized enrichment score|>1.0). Enrichment plots were plotted in R using the fgsea package version 1.26.0. Heatmaps were annotated in R with ComplexHeatmap package version 2.16.0^62^.

### Western blotting

DSRCT cells containing DOX-inducible shRNAs were treated with DOX (1 µg/ml) for 96 h. BER wild-type cells were siRNA-treated and incubated as described above. Whole cellular protein was extracted using RIPA buffer (SERVA Electrophoresis GmbH, Heidelberg, Germany) supplemented with 1X Halt™ Protease and Phosphatase Inhibitor Cocktail (Thermo Scientific). After quantitation by bicinchoninic acid assay (BCA) (Thermo Scientific), proteins were separated on a 10% SDS-PAGE gel at 100 V and blotted on PVDF membranes using the Trans-Blot Turbo Transfer System (BioRad). Membranes were incubated with mouse monoclonal anti-CACNA2D2 (1:1,000, sc-365911, Santa Cruz Biotechnology, Inc., Dallas, TX, USA), rabbit polyclonal anti-WT1 (1:500, sc-192, Santa Cruz), rabbit polyconal anti-EWSR1 (1:1000, #11910, Cell Signaling Technology Europe B.V. Leiden, The Netherlands) or rabbit monoclonal anti-GAPDH (1:1,000, #2118, Cell Signaling Technology Europe B.V. Leiden, The Netherlands). Then, membranes were incubated with horseradish peroxidase (HRP) coupled anti-rabbit IgG (1:5,000, sc-2357, Santa Cruz) or anti-mouse IgG (1:5,000, A9044, Sigma-Aldrich). Proteins were detected using chemiluminescence and Immobilon Western HRP Substrat (Sigma-Aldrich).

### In vivo experiments

For subcutaneous experiments, 100 μl of 1×10^6^ DSRCT cells harboring a shRNA against *WT1*-CDS or *WT1*-UTR were resuspended in PBS (Thermo Scientific) and mixed with Geltrex™ LDEV-Free, hESC-Qualified, Reduced Growth Factor Basement Membrane Matrix (Matrigel) (Thermo Scientific) in 1:1 proportion, and slowly injected subcutaneously in the right flank of 10–12 weeks old NOD/scid/gamma (NSG) mice as previously described^63^. The bodyweight of the mice was measured twice a week. Tumor diameters were measured every second day with a caliper and tumor volume was calculated by the formula L×l^2^/2, where L is the length and l the width of the tumor. When the tumors reached an average volume of 80 mm^3^, mice were randomized into two groups. 96 h prior to the pre-determined end of the experiment, mice were treated with either 2 mg/ml DOX (Thermo Scientific) dissolved in drinking water containing 5% sucrose (Carl Roth) to induce an in vivo knockdown (KD) of WT1 (DOX(+)), or 5% sucrose (control, DOX(-)). At the experimental endpoint or if other humane endpoints were reached before (body weight loss of 20%, apathy, piloerection, self-isolation, aggression as a sign of pain, self-mutilation, motor abnormalities such as a hunched back and reduced motor activity, as well as any other unphysiological or abnormal body posture, breathing difficulties, maximum tumor size of 15 mm in any direction, or an ulcerating tumor), mice were sacrificed by cervical dislocation. For orthotopic experiments, 100 μl of 1×10^6^ DSRCT cells harboring a shRNA against WT1-CDS or WT1-UTR in a 1:1 mix of cells suspended in PBS (Thermo Scientific) with Geltrex™ LDEV-Free, hESC-Qualified, Reduced Growth Factor Basement Membrane Matrix (Matrigel) (Thermo Scientific) were slowly injected in the peritoneum of 10–12 weeks old NOD/scid/gamma (NSG) mice. Seven-weeks after injection, mice were randomized in two groups and treated with either 2 mg/ml DOX (Thermo Scientific) dissolved in drinking water containing 5% sucrose (Carl Roth) to induce an in vivo knockdown of WT1 (DOX (+)), or 5% sucrose (control, DOX (−)) for 96 h. At the experimental endpoint or if the aforementioned humane endpoints were reached before, mice were sacrificed by cervical dislocation. Then, subcutaneous or orthotopically xenografted tumors were extracted. A section of each tumor was resected, snap frozen and reserved for RNA or protein extraction to confirm WT1 knockdown efficiency. Another section of each tumor was used for preparation of a single-cell suspension for scRNA-seq. The remaining tumor masses were fixed in 4%-formalin (Carl Roth) and paraffin-embedded (FFPE) for (immuno)histological analysis. Experiments were approved by the government of North Baden and conducted in accordance with ARRIVE guidelines and recommendations of the European Community (86/609/EEC) and UKCCCR (guidelines for the welfare and use of animals in cancer research).

### Human samples and ethics approval

All samples analyzed in this study had undergone rigorous morphological examination by reference pathologists and cancer-type specific molecular testing including identification of EWSR1::WT1 for DSRCTs, whenever possible. DSRCT morphological mimics included alveolar soft part sarcoma (ASPS), Ewing sarcoma, ganglioneuroblastoma, leiomyosarcoma, liposarcoma, malignant fibrous histiocytoma, nephroblastoma, neuroblastoma, osteosarcoma, rhabdomyosarcoma, synovial sarcoma, gastrointestinal stromal tumor (GIST), mesothelioma, hepatoblastoma, and unspecified small round cell sarcomas. Open slides or tissue-microarrays (TMAs) from human FFPE or cryopreserved tissue samples were retrieved from the archives of the Institute of Pathology of the LMU Munich, the Charité Berlin, The Biobank of the Hospital Universitario Virgen del Rocío of Seville, the Hospital Gustave Roussy (Villejuif), the Bone Tumor Reference Center at the University of Basel, the University of Essen, the Cooperative Weichteilsarkom Studiengruppe (CWS) study center, the Klinikum Stuttgart (ethics committee from the Medical Faculty of the Eberhard-Karls University and University Hospital of Tübingen, approval no. 207/2022BO2), the Radboud University Medical Center, the Pathology Institute of the LMU Munich (approval no. 550-16 UE), and the University of Heidelberg (approval no. S-211/2021).

### Immunohistochemistry (IHC) and immunoreactivity scoring

Paraffin-embedded tissue sections (3–4 μm) were deparaffinized and rehydrated in distilled water. Antigen retrieval was performed using a steamer with Citrate Buffer pH 6.0 for 20 min at 98°C, followed by cooling to room temperature. After rinsing with Tris Buffered Saline with 0.05% Tween 20 (TBS-T, Sigma-Aldrich), sections were blocked with blocking solution BLOXALL (Vector Laboratories, Newark, CA, USA) for 15 min at RT. Subsequently, they were incubated with 2.5% Horse Serum (Vector Laboratories) for 25 min at RT to minimize non-specific binding. Following two washes with TBS-T, sections were incubated with the primary monoclonal antibody against CACNA2D2 (1:3,000, sc-365911, Santa Cruz) in antibody diluent (Agilent technologies, Santa Clara, CA, USA) for 2h at 37°C. After washing three times with TBS-T, sections were incubated with a secondary horseradish peroxidase (HRP)-coupled horse-anti-rabbit/mouse antibody (ImmPRESS HRP Universal PLUS Polymer Kit (Peroxidase, Horse Anti-Rabbit/Mouse IgG)) (Vector Laboratories) for 30 min at RT. After three washes with TBS-T, chromogen staining was performed using 3,3’-diaminobenzidine (DAB) for 10 min at RT. Sections were then rinsed in distilled water for 5 min followed by tap water for 10 min. Counterstaining was performed using Hemalum (Carl Roth) for 1 min, followed by rinsing in tap water for 10 min. Finally, sections were dehydrated, cleared, and mounted with aqueous mounting media (Sigma-Aldrich). Evaluation of CACNA2D2 immunoreactivity was carried out in analogy to scoring of hormone receptor Immune Reactive Score (IRS) ranging from 0–12. This modified IRS scoring scheme has been adapted to Ewing sarcoma and been described and validated previously^25,64–67^. The percentage of cells with expression of the given antigen was scored and classified in five grades (grade 0=0−19%, grade 1=20−39%, grade 2=40−59%, grade 3=60−79% and grade 4=80−100%). In addition, the intensity of marker immunoreactivity was determined (grade 0=none, grade 1=low, grade 2=moderate and grade 3=strong). The product of these two grades defined the final IRS.

### Automated sample preparation (autoSP3) for proteome profiling

Samples were prepared as previously described^68^ unless otherwise stated. In brief, cell pellets were resuspended in 75 µl SDS lysis buffer (4% SDS, 100 mM Ammonium bicarbonate pH 8.5). For cell lysis and protein extraction, samples were subjected to AFA-ultrasonication using LE220R-plus ultrasonicator (Covaris Ltd, UK). Subsequently, protein concentrations were determined using the BCA Protein Assay Kit (Pierce, Thermo Fisher) and 20 µg of protein per sample were used as a direct input for the autoSP3 protocol. The autoSP3 protocol including protein clean-up, reduction and alkylation (using 10 mM TCEP and 40 mM CAA at final concentration) and digestion (trypsin, enzyme:protein ratio of 1:20), were performed on the Bravo liquid handling system (Agilent Technologies) as previously described^69^ using the Paramagnetic beads for SP3 (Sera-Mag Speed Beads A and B) (Fisher Scientific). The resulting peptides were directly frozen at −80°C until MS acquisition.

### Mass spectrometry data acquisition and data processing

Equivalent amount of 200 ng peptides per sample were injected into the timsTOF Pro mass spectrometer (Bruker Daltonics) coupled with an Easy nLC 1200 system (Thermo Scientific) fitted with an analytical column (Aurora Column with CSI fitting, C18, 1.6 μm, 75 μm x 25 cm) (Ion Optics). The elution gradient was set to 80 min at a flow rate of 300 nl/min using solvent A (0.1% formic acid in ULCM grade water) and solvent B (0.1% formic acid in 80% acetonitrile and 19.9% ULCM grade water). Data was acquired in DIA-PASEF mode. The full scan MS spectra was set to mass range 100 to 1,700 m/z and 1/k0 range from 0.65 to 1.42 V*s/cm^2^ with 100 ms ramp time. The duty cycle was locked at 100%, the ion polarity was set to positive, and TIMS mode was enabled. Collision energy was set to 1/k0 range from 0.65 to 1.42 V*s/cm^2^. For the DIA scans, a custom isolation window pattern was optimized, covering the precursor range of 377 to 1,194 m/z, mobility range 1/k0 range from 0.67 to 1.39 V*s/cm^2^, and cycle time estimate of 1.58 s. The resulting raw files were processed with DIA-NN software (version 1.8.1)^70^ using the default settings unless stated otherwise. Trypsin/P was selected as digesting enzyme, maximum missed cleavages was set to 2. N-term M excision, C carbamidomethylation, and oxidation were set as fixed and variable modifications. Match-between-runs (MBR) function was allowed. The DIA-NN search was performed using the *H.sapiens* Uniprot database (reviewed only, downloaded on 30.03.2021), and in silico spectral library generated by FragPipe using the previously mentioned fasta file.

### UMAP clustering of methylation array data

Preprocessed beta values from methylation arrays from reference cases of the Sarcoma classifier^71^ were imported into R. Dimensionality of the data was reduced using Uniform Manifold Approximation and Projection (UMAP) with the umap R package version 0.2.10.0 ^72,73^. Parameters of UMAP were number of neighbors: 30, minimum distance: 0.1. Clustering results were visualized on the UMAP plot using color-coded labels for different entities using ggplot2 R package version 3.4.4^74^.

### Statistical analyses

If not otherwise specified, statistical data analysis was performed using PRISM 9 (GraphPad Software Inc., CA, USA) on the raw data. If not otherwise specified in the figure legends or the respective methods section, comparison of two groups in functional in vitro experiments was conducted using an unpaired two-sided Mann-Whitney test. Pearson correlation analysis was done in R using Stats package version 4.3.0 and visualized in correlation matrices plotted in R with the corrplot package version 0.92 package. If not otherwise specified in the figure legends, data are presented as box-dot plots with horizontal bars representing means and whiskers representing the standard error of the mean (SEM). Sample size for all in vitro experiments was chosen empirically. For in vivo experiments, the sample size was predetermined using power calculations with *β* = 0.8 and *α*<0.05 based on preliminary data and in compliance with the 3R system (replacement, reduction, refinement). *P*-values<0.05 were considered as statistically significant. If not otherwise specified in the figure legends, all *P*-values were estimated from nonparametric two-sided statistical tests.

### Data availability

All data supporting the findings of this study are available within the article and its supplementary information files or from the corresponding author upon reasonable request.

## RESULTS

### Integrative multi-omic analysis identifies CACNA2D2 as significantly overexpressed in DSRCT compared to morphological mimics

We hypothesized that exploring a specific clinically-relevant gap in our understanding of DSRCT would illuminate the missing scientific resources necessary to bridge it, while simultaneously highlighting the already existing datasets that could be further curated and exploited. Hence, we initially addressed the fundamental clinical challenge of guaranteeing a fast, inexpensive, and reliable diagnosis of DSRCT and employed it as a guide in our efforts to generate a comprehensive toolset for DSRCT research. To identify a highly specific diagnostic biomarker, we focused on genes overexpressed in DSRCT compared to other cancer entities of likely differential diagnostic relevance (morphological mimics), including SRCSs, such as Ewing sarcoma, *BCOR*-rearranged sarcomas, and other *EWSR1*-rearranged malignancies^1,50,75,76^. To that end, we first aggregated publicly available microarray RNA expression data from 20 morphological mimics and combined them with expression data of 32 DSRCT primary patient samples that had previously been exclusively used as a reference dataset for studies on Ewing sarcoma^51^. Stringent paired differential gene expression (DEG) analysis comparing the expression profile of DSRCT with that of each mimic entity for the 20,360 genes represented on the microarray platform (**Fig. 1A**), identified 23 genes displaying a significant overexpression in the DSRCT samples (log2 expression fold change (log2FC)>2.5 and adjusted *P*-value (*Padj*)<0.01) (**Fig. 1B**). To further refine the selection of candidate genes, we hypothesized that genes highly expressed across DSRCTs would be likely driven by the pathognomonic EWSR1::WT1 fusion. To test this hypothesis, we collected publicly available chromatin immunoprecipitation followed by sequencing (ChIP-seq) data from the JN-DSRCT-1 cell line and searched for high-confidence binding sites of EWSR1::WT1 (GEO accession code GSE156277)^14^. This analysis identified 2,065 genomic loci bound by EWSR1::WT1 (**Fig. 1B**). To further enrich these results, we explored the DSRCT proteome for the first time, bridging an important gap in the available datasets for DSRCT. For this, we established cell line models from JN-DSRCT-1 and another DSRCT cell line (SK-DSRCT2), which stably expressed DOX-inducible shRNAs targeting either the coding sequence (CDS) or the 3’ untranslated region (3’-UTR) of *EWSR1::WT1*^77,78^. After 96 h of DOX-treatment the subsequent KD of EWSR1::WT1 was confirmed by western blotting (**Suppl. Fig. 1**), and the cells’ whole proteome was analyzed by mass spectrometry. Next, we performed differential protein expression (DEP) analysis to identify proteins regulated by EWSR1::WT1. This analysis yielded 104 proteins being concurrently regulated in both cell lines, expressing either shRNAs against *WT1* (log2FC>1.0 and *Padj*<0.01) (**Fig. 1B, Suppl. Table 6**).

**Fig. 1.**
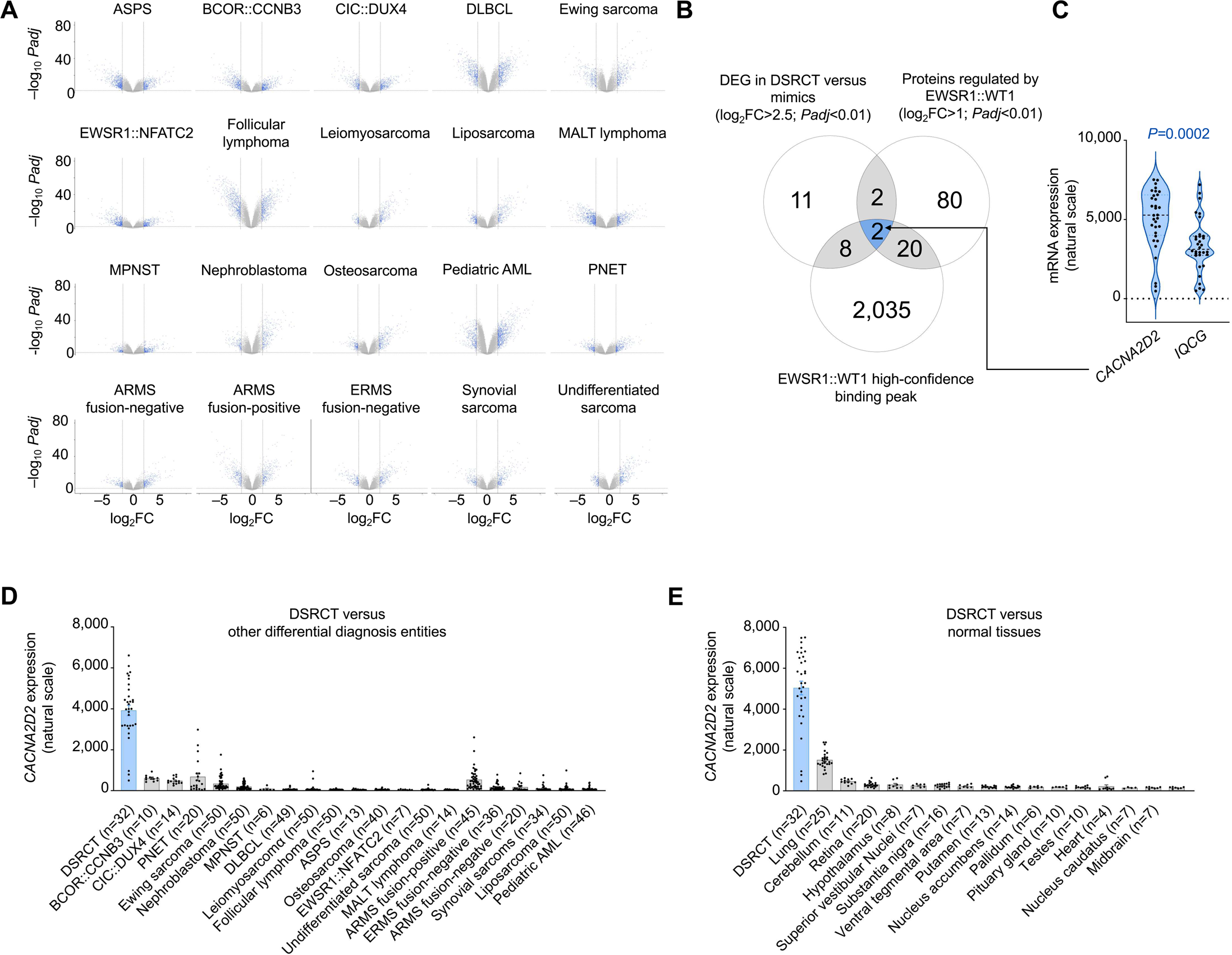
| Integrative multi-omic analysis identifies CACNA2D2 as significantly overexpressed in DSRCT compared to morphological mimics. **A.** Volcano plots depicting results of pairwise differential gene expression analysis of microarray mRNA gene expression profiles from primary DSRCT tumor samples and the indicated tumor entity. Blue dots represent genes with |log2FC|>2.5 and an *Padj*<0.01 (Benjamini-Hochberg corrected). Alveolar soft part sarcoma (ASPS); diffuse large B-cell lymphoma (DLBCL); mucosa-associated lymphoid tissue (MALT) lymphoma; malignant peripheral nerve sheath tumor (MPNST); acute myeloid leukaemia (AML); primitive neuroectodermal tumor (PNET), alveolar rhabdomyosarcoma (ARMS); embryonal rhabdomyosarcoma (ERMS). **B.** Venn diagram showing the overlap of genes significantly overexpressed (log2FC>2.5; *Padj*<0.01) in DSRCT compared to all other tumor entities as in **A** with genes or proteins potentially regulated by EWSR1::WT1 as determined by ChIP-seq and Mass spectrometry analyses. **C.** Violin plot showing *CACNA2D2* and *IQCG* mRNA expression in 32 DSRCT patient samples. Black dotted line indicates median. Blue dotted lines indicate quartiles. Unpaired two-sided Mann-Whitney test. **D.** *CACNA2D2* mRNA expression in 32 DSRCT patient samples and in 654 patient samples from 20 different entities. Horizontal bars represent mean expression levels, and whiskers SEM. The number of analysed samples is given in parentheses. **E.** *CACNA2D2* mRNA expression in 165 samples from the 15 different normal tissue types with the highest *CACNA2D2* expression as compared to 32 DSRCT patient samples. Horizontal bars represent mean expression levels, and whiskers SEM. The number of analysed samples is given in parentheses. Unpaired two-sided Mann-Whitney test.

The intersection of our DEG, ChIP-seq peaks, and DEP analyses revealed two proteins as highly promising diagnostic biomarkers in DSRCT: calcium voltage-gated channel auxiliary subunit alpha2delta 2 (CACNA2D2) and IQ motif containing G (IQCG) (**Fig. 1B**). A comparison of the expression of these two candidates in 32 DSRCT samples revealed that *CACNA2D2* expression was on average 150% higher than that of *IQCG* (*P*=0.0002) (**Fig. 1C**). Hence, we focused on exploring CACNA2D2 as a potential diagnostic marker for DSRCT.

To obtain first clues on the specificity of CACNA2D2 for DSRCT, we evaluated its mRNA expression level in DSRCT relative to morphological mimics. Notably, DSRCT exhibited a significantly higher (*P<*0.0001) mRNA expression of *CACNA2D2* than any other analyzed morphological mimic or pediatric tumor entity, thus emphasizing the potential of CACNA2D2 as a novel and highly specific biomarker for DSRCT (**Fig. 1D**). Further, since IHC evaluations are often performed on biopsies where tumor tissue occurs in proximity with surrounding normal tissues, and DSRCT patients usually present with metastatic disease to a wide variety of organs, discernibility of micrometastases from normal tissues plays an essential role in correct diagnostic workup^1^. Excitingly, analysis of publicly available gene expression data from 929 normal tissue samples derived from 71 tissue types^64^ in comparison with 32 DSRCT patient samples revealed that *CACNA2D2* was highly and significantly (*P<*0.0001) overexpressed in DSRCT compared to any other analyzed normal tissue (**Fig. 1E**). In fact, DSRCT showed on average a 3.3-fold higher expression of *CACNA2D2* as compared to the highest *CACNA2D2*-expressing normal tissue (lung) (**Fig. 1E**).

Collectively, the implementation of our integrative data analysis approach identified *CACNA2D2* as a significantly overexpressed protein in DSRCT compared to morphological mimics and normal tissues.

### *CACNA2D2* is directly regulated by an EWSR1::WT1-bound SE

Our initial ChIP-seq analysis suggested that *CACNA2D2* might be a direct downstream target of EWSR1::WT1 (**Fig. 1B**). To further characterize this putative interaction, we crossed ChIP-seq data of the *CACNA2D2* locus of two independently generated public datasets from JN-DSRCT-1 cell line (GEO accession codes GSE156277 and GSE212977) and observed that EWSR1::WT1 bound to the *CACNA2D2* locus concurrently with RNA polymerase II (**Fig. 2A**). This binding occurred approximately 5,000 bp upstream of exon 2 within the first intron of *CACNA2D2*, suggesting a potential regulatory role of EWSR1::WT1 through an enhancer interaction (**Fig. 2A**). To explore the regulation of *CACNA2D2* by EWSR1::WT1 in more detail, we conducted binding motif analysis of published EWSR1::WT1 genomic binding coordinates in JN-DSRCT-1 (ref. ^14^). For this, the top five de novo EWSR1::WT1 recognition motifs were employed to perform Find Individual Motif Occurrences (FIMO) analysis focusing on the upstream region of *CACNA2D2* comprising the first two exons and the first intron (chr3:50,513,549–50,541,675, hg19). Interestingly, the region containing EWSR1::WT1 ChIP-seq binding peaks at the *CACNA2D2* locus (chr3:50,517,880–50,521,290, hg19) displayed up to 17 EWSR1::WT1 de novo binding motifs in JN-DSRCT-1 (**Fig. 2A**). The additional presence of RNA polymerase II binding sites in proximity of the *CACNA2D2* promoter region was further suggestive of transcriptomic activity at *CACNA2D2* locus in DSRCT cells (**Fig. 2A**). Hence, we explored this regulation in JN-DSRCT-1 cell line and four primary DSRCT patient samples (termed DSRCT_1–DSRCT_4). Histone mark analysis using publicly available data on activating epigenetic modifications of Histone 3 residues (H3K9ac H3K4me3, H3K4me1, and H3K27ac) (GEO accession code GSE212977) revealed enrichment of H3K9ac and H3K4me3 signal at the *CACNA2D2* promoter region, indicating transcriptional activity at the transcription start site (**Fig. 2A**). Further, enriched H3K4me1 and H3K27ac marks at the *CACNA2D2* locus indicated the presence of a potential active enhancer site at the EWSR1::WT1 binding region (**Fig. 2A**). Analysis of the four primary DSRCT samples mirrored those results obtained in JN-DSRCT-1, underscoring its potential as a valuable cell model for exploration of DSRCT (**Fig. 2A**). In line with our previous findings, shRNA mediated KD of EWSR1::WT1 in JN-DSRCT-1 led to an almost complete loss of the EWSR1::WT1 signal and H3K27ac enhancer marks at the *CACNA2D2* locus (**Fig. 2A**).

**Fig. 2.**
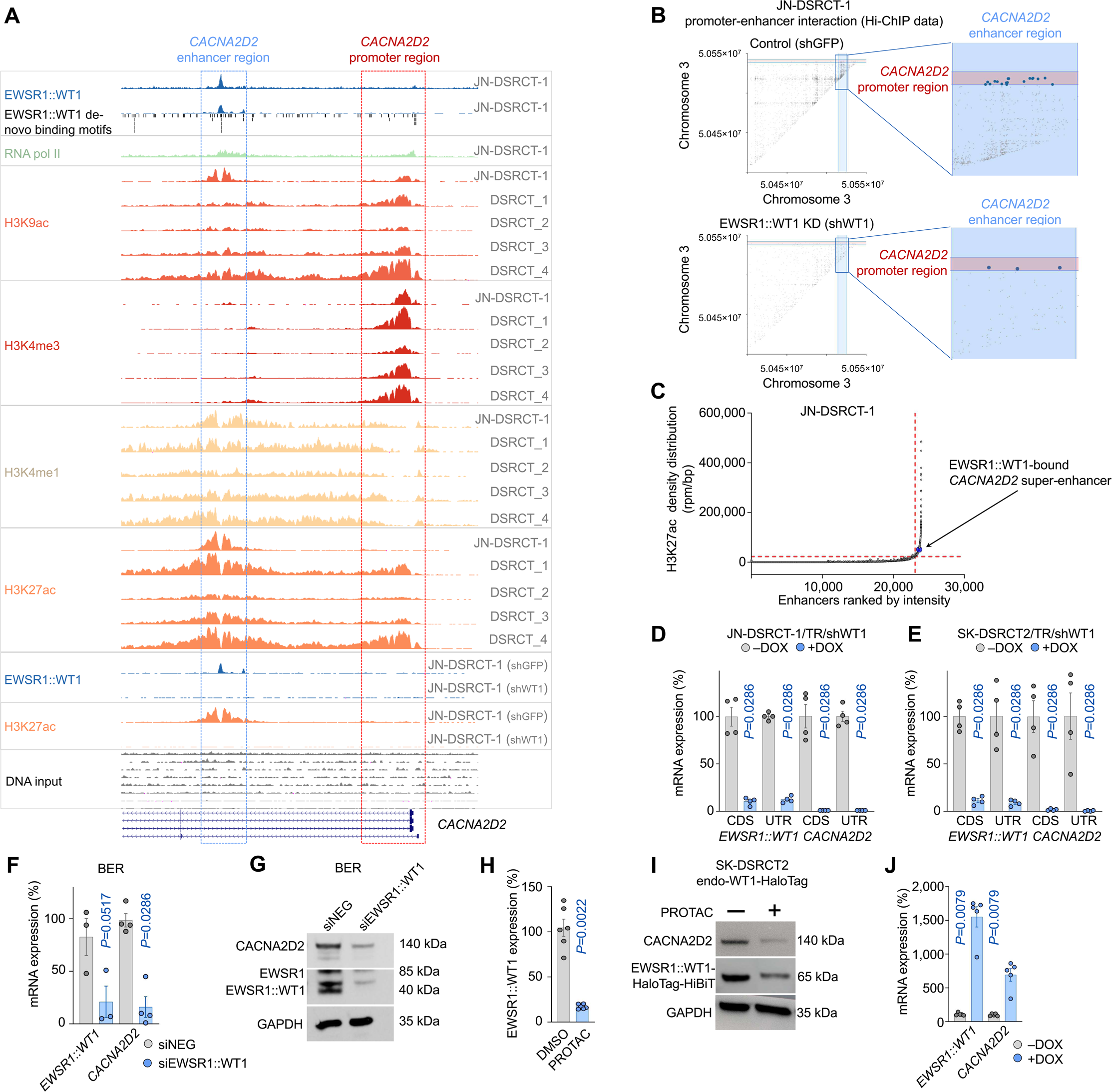
| *CACNA2D2* is directly regulated by a EWSR1::WT1-bound super-enhancer. **A.** EWSR1::WT1 de novo binding motifs and data of the epigenetic profile of *CACNA2D2* locus (chr3:50,506,517-50,548,707) from four primary DSRCT tumor samples (DSRCT_1-4), JN-DSRCT-1 wild-type, or a JN-DSRCT-1 cell line carrying a shRNA against GFP (control) or EWSR1::WT1. ChIP-seq profiles derived from GSE156277 and GSE212977 are depicted for EWSR1::WT1 (blue), RNA polymerase II (green), H3K4me3 (red), H3K4me1 (yellow), H3K27ac (orange), and H3K9ac (salmon). CACNA2D2 promoter and enhancer regions are highlighted with dotted squares. **B.** x-y scatterplot showing the spatial interactions between genomic loci in chromosome 3 captured by the Hi-ChIP experiment (GSE212978). The dot density reflects the frequency of interactions. Highlighted dots represent interactions of the *CACNA2D2* promoter and enhancer chromatin loci. The area colored in red indicates the *CACNA2D2* promoter region. The area colored in blue indicates the *CACNA2D2* enhancer region. **C**. x-y scatterplot showing H3K27ac signal density in reads per million mapped reads per base pair at active enhancer sites in JN-DSRCT-1 cell line ranked to their normalized intensity from low to high. Horizontal dashed red line indicates cut-off value of 22,119.5 for identification of super-enhancers (n=859 as indicated by vertical dashed red line) using ROSE algorithm. **D,E**. Bar plot showing relative mRNA expression levels of *EWSR1::WT1* and *CACNA2D2* as quantified by qRT-PCR in JN-DSRCT-1 (**D**) and SK-DSRCT2 (**E**) cell lines expressing a DOX-inducible shRNA mediated knock down of EWSR1::WT1. Cells were grown either with or without DOX for 96 h. Horizontal bars represent mean expression levels and whiskers SEM. n=4 biologically independent experiments. Unpaired two-sided Mann-Whitney test. **F.** Bar plot showing relative mRNA expression levels of *EWSR1::WT1* and *CACNA2D2* as quantified by qRT-PCR in BER DSRCT cell line transfected with siRNAs either targeting *EWSR1::WT1* (siEWSR1::WT1) or negative control (siNEG) for 48 h. Horizontal bars represent mean expression levels and whiskers SEM. n ζ 3 biologically independent experiments. Unpaired two-sided Mann-Whitney test. **G**. Western blot using antibodies against CACNA2D2, EWSR1(-WT1) and GAPDH (loading control) in BER cells transfected with siRNAs either targeting *EWSR1::WT1* (siEWSR1::WT1) or negative control (siNEG) 48 h. **H.** Bar plot showing relative expression levels of *EWSR1::WT1-HaloTag-HiBiT* transcript as quantified by luminescence in SK-DSRCT2-endo-WT1-HaloTag cells carrying a knock in of HaloTag-HiBiT tag to the endogenous locus of *EWSR1::WT1*. Cells were cultured with 1 μΜ PROTOAC or DMSO as control for 24 h. Horizontal bars represent mean expression levels and whiskers SEM. n=6 biologically independent experiments. Unpaired two-sided Mann-Whitney test. **I.** Western blot using antibodies against CACNA2D2 and (EWSR1::)WT1 in SK-DSRCT2-endo-WT1-HaloTag cell line carrying a knock in of HaloTag-HiBiT tag to the endogenous locus of EWSR1::WT1. Cells were cultured with 1 μΜ PROTAC or DMSO for 48 h. Loading control: GAPDH. **J.** Bar plot showing relative mRNA expression levels of *EWSR1::WT1* and *CACNA2D2* as quantified by qRT-PCR in MeT-5A-EWSR1::WT1^−KTS^ cell treated with or without DOX for 96 h. Horizontal bars represent mean expression levels and whiskers SEM. n=5 biologically independent experiments. Unpaired two-sided Mann-Whitney test.

In the context of these histone mark patterns, and given that other oncogenic fusion proteins, such as EWSR1::ETS and PAX::FOXO1, are described to regulate the expression of hallmark cancer genes via the formation of SEs^79,80^, and that EWSR1::WT1 mostly binds to intronic and intergenic regions in DSRCT^14^ such as the ones seen at the *CACNA2D2* locus in our data (**Fig. 2A**), we explored physical chromatin interaction of the putative *CACNA2D2* enhancer with the *CACNA2D2* promoter using H3K27ac Hi-ChIP data (GEO accession code GSE212976). This analysis revealed 19 high-quality chromatin interaction loops between the putative enhancer and the *CACNA2D2* promoter, mapping the EWSR1::WT1 binding site to the transcription start site of *CACNA2D2* (**Fig. 2B**). In line with these observations, shRNA mediated KD of *EWSR1::WT1* resulted in an 85% reduction of chromatin interaction at this locus (**Fig. 2B**). To further assess the potential role of *CACNA2D2* as an essential gene for DSRCT cell identity and function, we conducted SE analysis using ChIP-seq data on H3K27ac histone marks in JN-DSRCT-1 cell line. Notably, our analysis showed that the EWSR1::WT1-bound enhancer associated with *CACNA2D2* exhibited a characteristic SE H3K27ac profile of a so-called SE^33,81^ (**Fig. 2C**), which was lost upon EWSR1::WT1 KD (**Suppl. Table 7**), implying that *CACNA2D2* expression could be regulated by an EWSR1::WT1-bound SE (**Fig. 2C**).

To confirm the regulation of CACNA2D2 expression by EWSR1::WT1 using in vitro modeling (**Fig. 1B**, **Fig. 2A,B**), we next analyzed the mRNA and protein expression patterns of CACNA2D2 upon post-transcriptional and post-translational repression of EWSR1::WT1 in DSRCT cell line models. In a first step, we used the cell line models carrying each of the shRNAs (shWT1-CDS and shWT1-UTR) against *EWSR1::WT1* in JN-DSRCT-1 and SK-DSRCT2 and found that efficient KD of EWSR1::WT1 mRNA and protein (JN-DSRCT-1 average 12% remaining mRNA expression; SK-DSRCT2 average 10% remaining mRNA expression) results in a significant reduction of *CACNA2D2* expression in all tested cell line models (**Fig. 2D,E, Suppl. Fig. 1A**). To further validate these results, we employed a third DSRCT cell line, BER, which was transiently transfected with siRNAs targeting the *EWSR1::WT1* transcript or negative control. Again, we observed a significant reduction of CACNA2D2 mRNA and protein expression upon KD of EWSR1::WT1 using this alternative RNA interference method, which further underscores that EWSR1::WT1 expression is necessary for *CACNA2D2* upregulation (**Fig. 2F,G**). Consistent with these results, proteolysis-targeting chimera (PROTAC)-driven EWSR1::WT1 protein degradation in SK-DSRCT2 resulted in a significant reduction of EWSR1::WT1, which was accompanied by a substantial reduction of CACNA2D2 mRNA and protein expression (**Suppl. Fig. 2A, Fig. 2H,I**). Importantly, each of the three DSRCT cell lines expresses different EWSR1::WT1 isoforms, all commonly found in patient samples^82^ (**Suppl. Fig. 2B**). Hence, our data suggests that CACNA2D2 is consistently upregulated in DSRCTs regardless of the present EWSR1::WT1 isoform.

To further elucidate whether EWSR1::WT1 expression is not only necessary but also sufficient to drive *CACNA2D2*, we employed two heterologous mesothelial cell line models described to be the potential cell of origin of DSRCT (LP-9 and MeT-5A)^7,83^. For this, analysis of publicly available ChIP-seq data from MeT-5A cells expressing either wild-type WT1 or EWSR1::WT1 isoforms (-KTS, +KTS, or both -KTS/+KTS) revealed that mesothelial cells expressing the +KTS or -KTS/+KTS isoforms exhibited significant binding of EWSR1::WT1 to the putative *CACNA2D2* enhancer region (GEO accession code: GSE212977, **Suppl. Fig. 2C**). Although data on MeT-5A cells expressing only the -KTS isoform is virtually missing in this dataset, H3K27ac histone marks were enriched at the *CACNA2D2* promoter of MeT-5A cells expressing the -KTS, +KTS, or -KTS/+KTS isoforms, suggesting that both isoforms alone could drive *CACNA2D2* expression in EWSR1::WT1-expressing MeT-5A cells (**Suppl. Fig. 2C**). To explore this possibility, we ectopically overexpressed *EWSR1::WT1*^−KTS^ in MeT-5A cells and excitingly confirmed that *EWSR1::WT1^−KTS^*was also sufficient to drive significant *CACNA2D2* expression (**Fig. 2J**). Of note, isoforms of wild type WT1 were not bound to the CACNA2D2 enhancer region, and an active enhancer site could not be detected, suggesting that CACNA2D2 regulation was a distinct feature of EWSR1::WT1-mediated oncogenic transformation (**Suppl. Fig. 2C**). To confirm the potential role of CACNA2D2 as a mediator of the potential transformation of MeT-5A cells, we performed SE analysis which strikingly showed that upon expression of EWSR1::WT1^−KTS+KTS^ the *CACNA2D2* enhancer bound by the fusion protein became a SE, hinting to a potential important role of CACNA2D2 in DSRCT (**Suppl. Fig. 2D**).

To further validate our findings in mesothelial cells and assess the transcriptional implications of EWSR1::WT1 binding to the *CACNA2D2* enhancer, we conducted DEG analysis using two independent publicly available RNA-seq datasets (GEO accession codes: GSE212976 and GSE252051) obtained from two different human mesothelial cell lines (MeT-5A and LP-9) expressing either -KTS, +KTS, or both EWSR1::WT1 isoforms. Our comprehensive data analysis revealed that regardless of the isoform, EWSR1::WT1 significantly drove *CACNA2D2* expression in both the MeT-5A and LP-9 cell lines (log2FCs ranging from 2.7–5.9, *Padj*<0.05) (**Suppl. Fig. 2E**).

Collectively, these findings revealed that *CACNA2D2* is a direct SE-driven downstream target of EWSR1::WT1 and that EWSR1::WT1 expression is necessary and sufficient to drive *CACNA2D2* expression.

### CACNA2D2 is a key component of the EWSR1::WT1 oncogenic signature

To continue our quest to generate a comprehensive toolset for DSRCT research, we further explored whether *CACNA2D2* could serve as a surrogate indicator of oncogenic *EWSR1::WT1* transformation, and hypothesized that *CACNA2D2*-associated gene expression patterns would overlap with those resulting from EWSR1::WT1 expression in DSRCT. Thus, as a first step, we performed a correlation analysis using microarray gene expression data from 32 DSRCT patient samples, revealing 2,873 coexpressed and 2,627 anti-coexpressed genes with *CACNA2D2* (|r|>0.3; *P*<0.05). These genes were used to define the *CACNA2D2* gene set for subsequent fast gene set enrichment analysis (fGSEA) (**Suppl. Table 8, Fig. 3A**). Additionally, genes with the highest ranking *CACNA2D2* correlation values (*n*=100) were selected to define the CACNA2D2 signature (**Suppl. Table 9, Fig. 3A**). Then, an EWSR1::WT1 signature was computed by first performing DEG analysis of microarray expression data obtained from in vivo subcutaneous xenograft material from two DSRCT cell lines transduced with two different shRNAs against the CDS and 3’UTR region of *WT1* (**Fig. 3A, Suppl. Table 9**). In addition, DEG analysis of publicly available RNA-seq data derived from the DSRCT cell line BER transfected with siRNAs targeting *EWSR1::WT1* was performed^14^ (**Fig. 3A**). By analyzing the overlap between these two resulting datasets we identified 87 genes significantly upregulated (log2FC>1.0, *Padj*<0.01) by EWSR1::WT1, representing the core *EWSR1::WT1* transcriptional signature in three DSRCT cell lines (**Suppl. Table 9, Fig. 3A**). Excitingly, fGSEA using the *CACNA2D2* gene set ranked using correlation coefficients as input showed a highly significant (*Padj*=3.4×10^−9^) and strong positive enrichment for the EWSR1::WT1 signature (NESEWSR1::WT1=3.6).

**Fig. 3.**
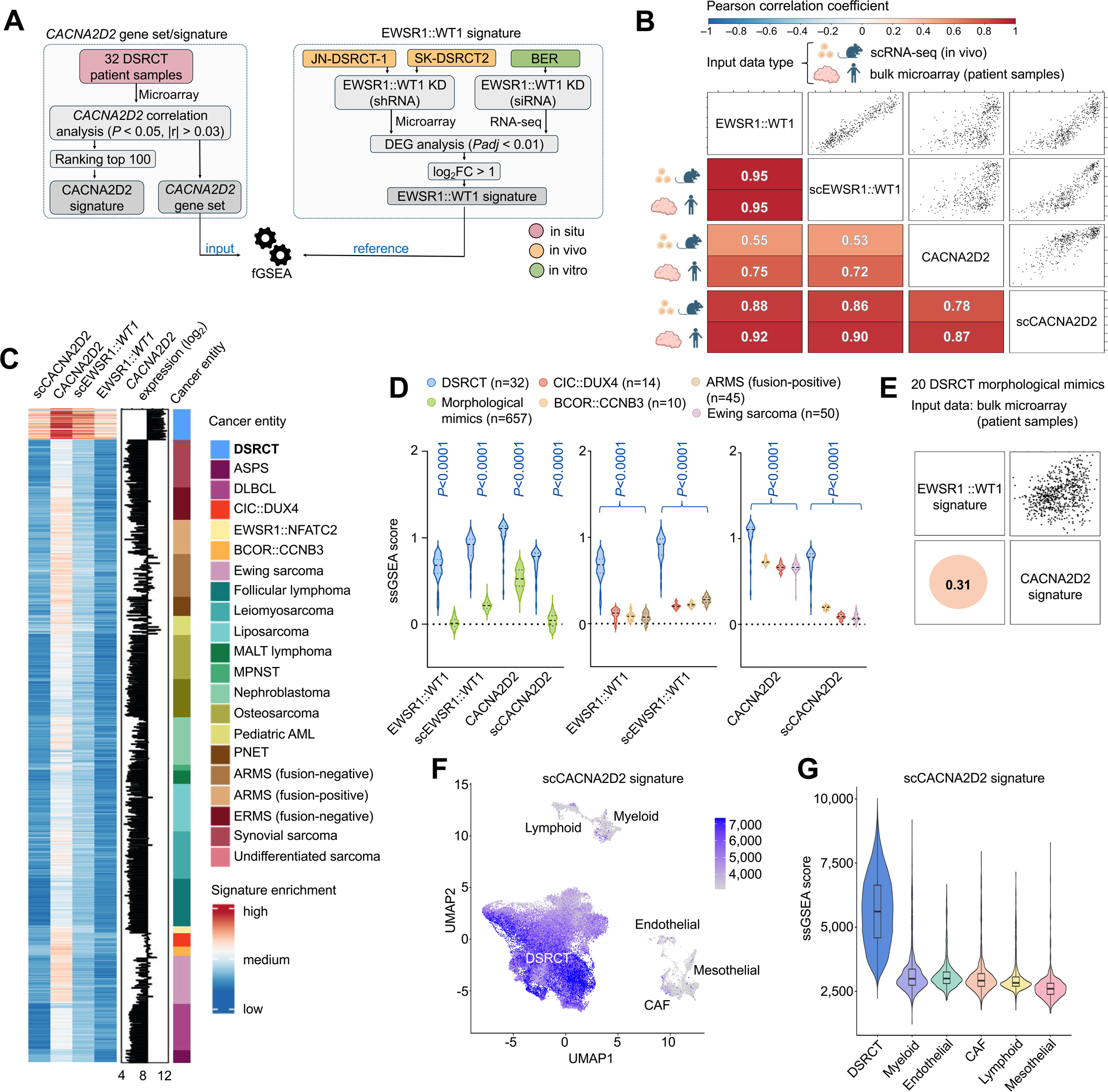
| CACNA2D2 is a key component of the EWSR1::WT1 oncogenic signature. **A.** Diagram depicting the workflow for establishment of the CACNA2D2 signature and *CACNA2D2* gene set, and EWSR1::WT1 signature and subsequent fGSEA using Pearson correlation and DEG analysis respectively. Color depicts the origin of the samples (in situ, in vivo, in vitro). **B**. x-y-scatter plots illustrating correlation of ssGSEA enrichment scores between (sc)EWSR1::WT1 and (sc)CACNA2D2 signatures of orthotopically xenografted single DSRCT cells. Displayed numbers and colors in the correlation matrix represent Pearson correlation values for each signature enrichment in single cells from orthotopically xenografted DSRCT (top) or 32 bulk DSRCT patient samples (bottom). **C**. Heatmap depicting ssGSEA signature enrichment scores for (sc)EWSR1::WT1 and (sc)CACNA2D2 signatures and log2 expression values for *CACNA2D2* across 32 DSRCT patient samples and 20 additional cancer entities of DSRCT differential diagnosis. Alveolar soft part sarcoma (ASPS); diffuse large B-cell lymphoma (DLBCL); mucosa-associated lymphoid tissue (MALT) lymphoma; malignant peripheral nerve sheath tumor (MPNST); acute myeloid leukaemia (AML); primitive neuroectodermal tumor (PNET), alveolar rhabdomyosarcoma (ARMS); embryonal rhabdomyosarcoma (ERMS). **D**. Left: violin plot comparing ssGSEA scores for (sc)EWSR1::WT1 and (sc)CACNA2D2 signatures in 32 DSRCT patient samples versus all other DSRCT-morphological mimics analyzed in **C**. Middle: violin plot comparing ssGSEA enrichment scores for (sc)EWSR1::WT1 in 32 DSRCT patient samples as compared to the top three scoring entities: BCOR::CCNB3, CIC::DUX4, fusion-positive RMS. Right: violin plot comparing ssGSEA enrichment scores for the (sc)CACNA2D2 signature in 32 DSRCT patient samples as compared to the top three scoring entities: BCOR::CCNB3, CIC::DUX4, and Ewing sarcoma. Dotted black lines represents the median and dotted blue lines quartiles. Unpaired two-sided Mann-Whitney test. **E**. Correlation matrix showing ssGSEA enrichment scores of EWSR1::WT1 and CACNA2D2 signatures across all 20 cancer entities of DSRCT differential diagnosis. **F**. UMAP plot of scRNA-seq analysis of merged and integrated data from eleven samples, comprising tumor-derived and normal cells from four DSRCT patients (GSE263523), using the scCACNA2D2 signature. Color gradient indicates the ssGSEA score for scCACNA2D2 signature enrichment. CAFs: Cancer-associated fibroblasts. **G**. Violin plot depicting scCACNA2D2 signature scores for DSRCT and other normal cell types included in the patient samples. Horizontal bar represents median. Cancer-associated fibroblasts (CAFs).

To further illuminate the likely interplay between *CACNA2D2* and *EWSR1::WT1,* we next conducted single sample gene set enrichment analysis (ssGSEA) of microarray expression data from 32 DSRCT patient samples. In line with our previous results, this analysis confirmed that the EWSR1::WT1 signature enrichment scores significantly and highly correlated with that of the CACNA2D2 signature (r=0.75), further highlighting a transcriptional interconnection between *CACNA2D2* and *EWSR1::WT1* in situ.

To explore whether these observations could be extrapolated to other DSRCT models at single-cell (sc) resolution, we generated orthotopically-derived tumors using the two DSRCT cell lines with conditional DOX-induced KD of EWSR1::WT1 in the peritoneum of NSG mice and treated them with either DOX or sucrose (control) for 96 h prior to tumor extraction. scRNA-seq of the obtained primary (n=221) and metastatic (n=221) tumor cells was performed. First, the single-cell-data-derived scCACNA2D2 gene signature was generated in analogy to the one derived from bulk sequencing by conducting correlation analyses of the scRNA-seq data to identify the top 100 highest *CACNA2D2-*correlating genes (**Suppl. Table 9**). To obtain the single-cell-data-derived scEWSR1::WT1 signature, we conducted DEG analysis of the same DOX or control tumor cells and defined significantly EWSR1::WT1-upregulated genes (log2FC>0.25, *Padj*<5×10^−5^, expression in >10% of all DOX-treated and control cells) across both cell lines expressing either shRNA against *EWSR1::WT1* (**Suppl. Table 9**). Next, we performed ssGSEA of our single-cell data which strikingly showed an almost perfect and highly significant correlation between enrichment scores of our bulk- and single-cell-derived EWSR1::WT1 signatures (r=0.95, **Fig. 3B**). Remarkably, both ssGSEA scores for the bulk- and single-cell-derived EWSR1::WT1 signatures exhibited high and significant correlations with scores for the bulk- and single-cell-derived CACNA2D2 signatures (r_EWSR1::WT1;scCACNA2D2_=0.88; r_scEWSR1::WT1;scCACNA2D2_=0.86; r_scEWSR1::WT1;CACNA2D2_=0.53) (**Fig. 3B**). Notably, analyses of single DSRCT cells extracted exclusively from metastatic sites showed a similar correlation strength between enrichment scores of our bulk- and single-cell-derived EWSR1::WT1 and CACNA2D2 signatures, implying that CACNA2D2*-*associated genes are a characteristic feature of metastasized DSRCT cells (**Suppl. Fig. 3A**).

To collectively validate these observations in situ, ssGSEA analysis of the bulk microarray expression data from 32 DSRCT patient samples was performed and consistently revealed a high correlation between each of the enrichment scores of both bulk- and single-cell-derived EWSR1::WT1 and CACNA2D2 signatures (**Fig. 3B**). This suggests that our generated gene signatures derived from both bulk- and single-cell data reliably capture the transcriptional interplay of *CACNA2D2* and *EWSR1::WT1* in orthotopically xenografted and in patient-derived tumors (**Fig. 3B**).

As anticipated, both bulk- and single-cell-derived EWSR1::WT1 signatures exhibited a notable overlap in gene composition (n=48 genes, comprising 55% of the bulk signature) further confirming their robustness (**Suppl. Fig. 3B**). Additionally, single-cell and bulk-derived EWSR1::WT1 and CACNA2D2 signatures presented an overlap of 69 genes, which emphasizes the tight regulatory connection between *EWSR1::WT1* and *CACNA2D2* expression (**Suppl. Fig. 3C**). To assess whether the observed high correlation between EWSR1::WT1 and CACNA2D2 signatures was primarily influenced by their shared gene pool, we conducted the previous ssGSEA analysis using single-cell- and bulk-derived ^exclusive^EWSR1::WT1 and ^exclusive^CACNA2D2 signatures, which excluded all genes shared by each CACNA2D2 and EWSR1::WT1 signature (**Suppl. Fig. 3C**). Strikingly, similarly high correlations between the different gene signatures were calculated (**Suppl. Fig. 3D**) suggesting that the interplay between EWSR1::WT1 and CACNA2D2 signature enrichment is not solely dependent on their shared gene pool, but rather involves both direct and indirect regulatory networks governed by EWSR1::WT1.

To delineate the specificity of the interaction of *CACNA2D2* and *EWSR1::WT1* expression patterns in DSRCT, we iterated ssGSEA using our newly generated bulk- and single-cell-derived EWSR1::WT1 and CACNA2D2 signatures of microarray expression data from 20 DSRCT morphological mimics of differential diagnosis (**Fig. 3C**). This analysis showed significantly lower enrichment scores in non-DSRCT analyzed tumor types for both signatures when compared to DSRCT samples (**Fig. 3D**). Additionally, correlation between these signatures was significantly lower for all analyzed DSRCT morphological mimics when compared to DSRCT (**Fig. 3E**). These results further emphasized the high specificity of the CACNA2D2 and EWSR1::WT1 interplay in DSRCT.

Finally, to test the strength of this newly defined CACNA2D2 signature in single-cell patient material that included non-tumor cells in combination with DSRCT cells, we used publicly available scRNA-seq data from four DSRCT patients (n=11 samples)^15^. Notably, both CACNA2D2 signatures (bulk and single-cell) precisely identified the cluster of DSRCT cells in all samples analyzed (**Fig. 3F, Suppl. Fig. 3E**). These results complemented the cell type annotation algorithms used to predict normal cell types in these tumors^15^, which all presented low enrichment of both CACNA2D2 signatures (**Fig. 3G, Suppl. Fig. 3F**).

In sum, the interplay between EWSR1::WT1- and CACNA2D2 signatures emerges as a distinctive hallmark of DSRCT, and as such, the CACNA2D2 signature can serve as a new tool to determine DSRCT cell identity.

### High CACNA2D2 protein expression is a diagnostic hallmark of DSRCT

Since our analyses highlighted a robust and specific interplay of *CACNA2D2* with *EWSR1::WT1* (**Figs. 1–3**), we further hypothesized that CACNA2D2 could be a specific diagnostic marker for DSRCT. First, to explore if the epigenetic status of *CACNA2D2* was sufficient to distinguish DSRCT from other tumor entities, we used publicly available DNA methylation array data from the Sarcoma Classifier^71^ and analyzed all 37 *CACNA2D2*-associated CpG sites. Strikingly, UMAP dimension reduction using exclusively these CpG sites of 24 DSRCT patient samples in comparison with 192 tumor samples from 13 other differential diagnostic sarcoma entities showed tight and distinct clustering of all DSRCT samples (**Fig. 4A**). Our analysis of the mean methylation levels of these *CACNA2D2*-associated CpG sites revealed a significant (*P*=0.0007) and specific hypomethylation in DSRCT patient samples compared to other differential diagnostic sarcomas, suggesting a specific epigenetic regulatory mechanism unique to DSRCT. (**Suppl. Fig. 4A**). To further define the specificity of *CACNA2D2*-driven DSRCT clustering as compared to other previously described EWSR1::WT1-regulated genes including *CCND1*, *NTRK3, BAIAP3, BLK, ENT4, FGFR4, JAK3, PDGFA,* and *IGF2*^5,7–9,11–14^, UMAP analyses of their corresponding CpG sites were performed. Excitingly, unlike the analysis of *CACNA2D2*-specific CpG sites, DSRCT clustering according to any other studied gene-specific CpG sites was not distinct (**Fig. 4A, Suppl. Fig. 4B**). Of note, UMAP analysis of CpG sites of *IQCG* (**Fig. 1C**) also failed to show clustering of DSRCT samples (**Suppl. Fig. 4B**). This data implies that *CACNA2D2*-associated methylation signature is a distinct and specific feature of DSRCT.

**Fig. 4.**
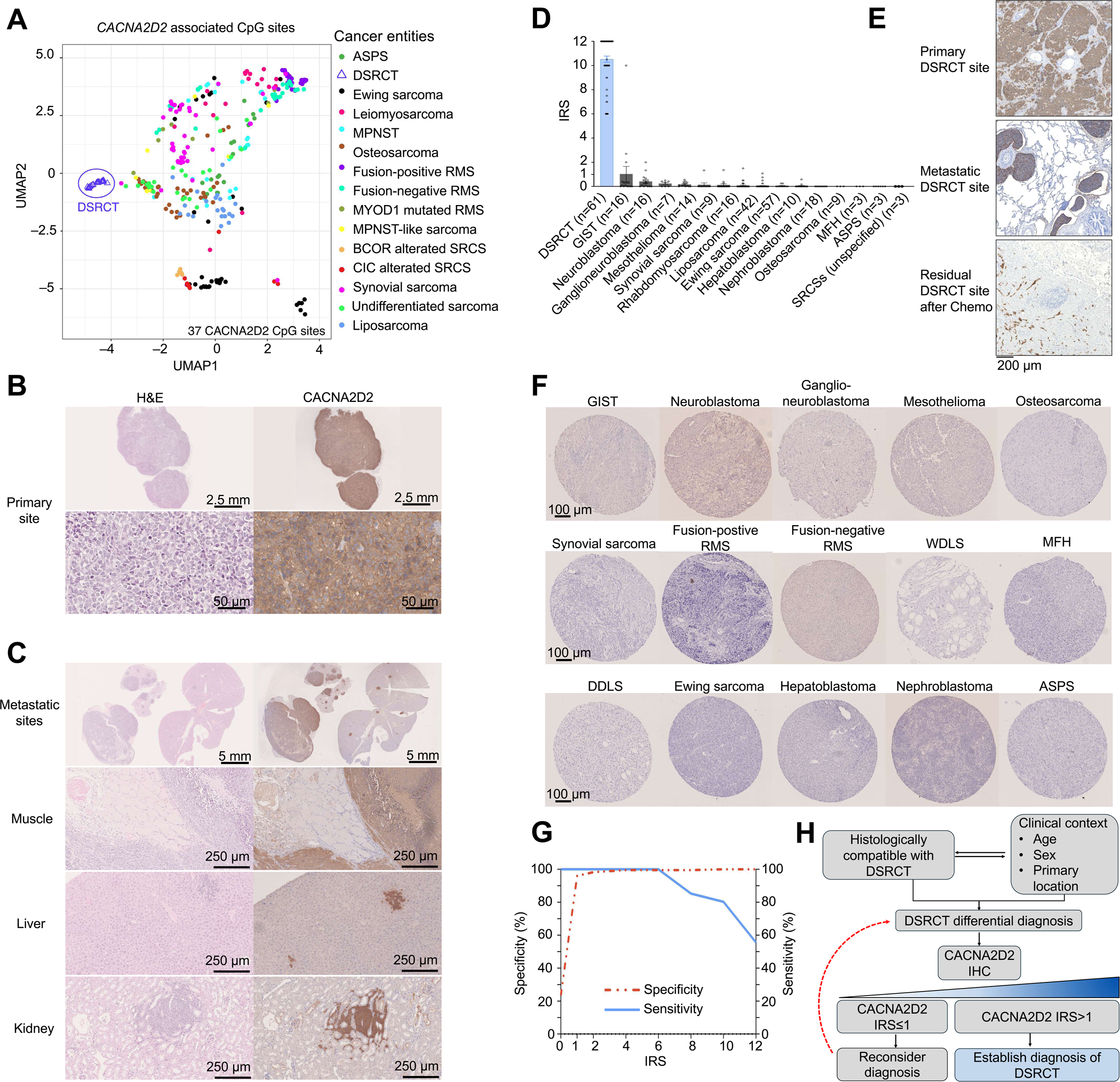
| High expression of CACNA2D2 is sufficient to robustly diagnose DSRCT by IHC. **A.** UMAP plot of 14 sarcoma entities, including DSRCT, according to their *CACNA2D2* methylation status. **B**. Representative histological images of CACNA2D2 IHC in a DSRCT murine orthotopic xenograft. DAB (brown chromogen) was used. **C**. Representative histological images of CACNA2D2 IHC in micro-metastases to inner organs (muscle, liver and kidney) from DSRCT murine orthotopic xenografts. A brown chromogen was used. **D**. Bar plot showing the individual IRS scores for CACNA2D2 staining in DSRCT and other morphological mimics. The number of analyzed samples is given in parentheses. Bars represent mean IRS values, whiskers indicate SEM. DSRCT samples are highlighted in blue color. Unpaired two-sided Mann-Whitney test. **E**. Top: Representative histological image of CACNA2D2 IHC in DSRCT primary tumor tissue. Middle: Representative histological image of CACNA2D2 IHC in a lung metastasis of DSRCT. Bottom: Representative histological image of CACNA2D2 IHC in a tumor sample with residual DSRCT cells after treatment. A brown chromogen was used. **F**. Representative histological images of CACNA2D2 IHC in TMA cores from different tumor types. A brown chromogen was used. **G**. Graph shows sensitivity (blue solid line) and specificity (red dashed line) of CACNA2D2 IHC. Both metrics are expressed as percentages. **H.** Diagram depicting this study’s proposed workflow for establishing robust diagnosis of DSRCT.

Next, to expand the translational implication of our findings *in vivo*, we optimized an IHC staining protocol of CACNA2D2 on our DSRCT cell line xenografts and obtained consistent and robust positive membranous or cytoplasmatic staining of entire tumor sections (**Fig. 4B**). Importantly, when assessing metastatic spread of these tumors to different organs in vivo, CACNA2D2 staining clearly and specifically detected DSRCT cells with high sensitivity, even uncovering small micrometastases in muscle, liver, or kidney tissue (**Fig. 4C**). This underscores the potential of CACNA2D2 staining as a reliable tool for both early detection of metastatic lesions, and assessment of microscopic residual disease, which is typically still present after chemotherapy and complete cytoreductive treatment in DSRCT patients^84^.

Finally, to prove whether the DSRCT-specific staining pattern of CACNA2D2 could be observed in primary patient samples and thus be employed as a DSRCT marker, we assembled the largest collection of fresh-frozen or FFPE DSRCT patient samples analyzed to date (n=61) comprising primary, metastatic and post-treatment samples. Further, we complemented this cohort with 229 patient samples from 15 different DSRCT morphological mimics of potential differential diagnostic relevance (**Suppl. Table 10**). We then assessed this sample collection using our optimized IHC staining protocol for CACNA2D2. Immunoreactivity scores (IRS) were determined in analogy to the Remmele and Stegner scoring system^85^ as previously employed in the DSRCT morphological mimic Ewing sarcoma^25,50,65,67,86,87^ (IRS range from 0–12). DSRCT tumor sections exhibited the highest IRS for CACNA2D2 when compared to all other analyzed cancer entities (IRS_mean_=10.5, *P*<0.0001) with remarkably high specificity, which reached 99% when applying the cutoff IRS>1 (**Fig. 4D–G**). Furthermore, the sensitivity of CACNA2D2 staining was 100% when applying an IRS cutoff of ≤6, implying that DSRCT samples generally show a strong positive staining for CACNA2D2 (**Fig. 4G**).

Thus, we propose the following workflow to establish a diagnosis of DSRCT (**Fig. 4H**): In the case of clinically suspected and histologically-compatible DSRCT differential diagnosis, a biopsy should be first stained for CACNA2D2. In the case of a CACNA2D2 staining with an IRS≤1, the potential DSRCT diagnosis should be reconsidered, and additional confirmatory molecular diagnostic procedures (such as FISH, qRT-PCR, and/or next-generation sequencing), if available, may be performed to further delineate the specific diagnosis. Conversely, if higher (membranous) expression of CACNA2D2 (defined as IRS>1) is histologically evaluated, a diagnosis of DSRCT may be established.

All in all, our initial clinically-relevant question on the lack of a specific DSRCT biomarker guided a comprehensive effort, which resulted in the generation of an extensive toolset for DSRCT research (**Suppl. Fig. 4C**) and a validated blueprint for how such resources could be harnessed in other cancer entities.

## DISCUSSION

Despite its tremendous unmet clinical need, there has been a notable lack of comprehensive studies dedicated to unraveling the molecular intricacies of DSRCT as compared to other rare malignancies^88^. The resulting challenges in understanding DSRCT are exacerbated and reflected by the lack of clinical trials or trial groups dedicated to this disease, ultimately leading to limited systematic collection and analysis of DSRCT biomaterial^1^. Efforts to profile the limited available DSRCT cell line models in vitro including post-transcriptional knockdown models of the oncogenic translocation^5,7,15,18^ and knockdowns of selected potential oncogenic key regulators in DSRCT^10,17,83^ have only recently been described. This gap underscores the difficulty in improving diagnostic and therapeutic approaches for DSRCT^1^.

To address this deficiency, our study provides a comprehensive multi-model approach to DSRCT from a multi-omic perspective. By characterizing for the first time both the proteome and single-cell transcriptome of DSRCT cell lines and expanding additional novel analyses to subcutaneous and orthotopically xenografted tumors, we provide a new toolset for an exhaustive perspective on this rare disease.

In addition, we have successfully established a robust pipeline leveraging multi-omic analyses and integrating a wide array of published and newly generated datasets, which have not been previously utilized for such purposes in DSRCT research. The data encompasses various omic-layers, allowing a holistic analysis of the molecular underpinnings of DSRCT, now also including a pipeline for SE analysis of the DSRCT epigenome, and that of mesothelial cells as its potential cell of origin. In addition, given that SEs are known to contribute to cell identity and are often found close to genes having cell type-specific function, our results demonstrate that DSRCT cell identity could be substantially determined by *CACNA2D2*^33,81^.

The newly generated datasets not only fill a significant gap but also open avenues for further exploration and potential diagnostic advances. Building on our multi-omic approach, we also established gene signatures highly specific to DSRCT encompassing a core set of genes — termed the CACNA2D2 signature— that are consistently dysregulated in DSRCT and reflect the fusion expression pathognomonic for DSRCT. The CACNA2D2 signature was rigorously validated using data from both in vivo and in situ materials, highlighting its robustness in defining DSRCTs.

Clinically, the lack of specific IHC markers poses an additional diagnostic challenge since a reliable diagnosis requires molecular testing. In this context, diagnosing DSRCT poses difficulties arising from its rarity and ambiguous histopathological features as it can present as neither desmoplastic nor be composed of (nests of) small or round tumor cells^3^. In fact, these tumors may be easily misdiagnosed as other reminiscent entities within the spectrum of morphological mimics, including small round cell sarcomas (SRCSs) or neuroendocrine tumors^75,89^. Best attempts to diagnose DSRCT via IHC have relied on the co-occurrence of multiple immunophenotypical stains, such as the combination of positive staining for EMA, Desmin and NSE^90^. In addition, the detection of C-terminal WT1, presents several challenges, including: a) atypical staining patterns may arise due to the expression of full-length WT1 or rare isoforms of EWSR1::WT1, requiring a deeper understanding of anti-WT1 antibody epitopes and their correlation with clinical and genetic findings^91^; b) all WT1 C-terminal antibodies used for DSRCT identification have been discontinued; and c) high WT1 expression is also seen in other cancers^92,93^. If available, these IHC approaches have been successfully complemented by detecting the *EWSR1::WT1* fusion at the DNA or RNA level using techniques such as FISH, RT-PCR or NGS^1^. However, these methods are more expensive and time-consuming than IHC, whereas a swift and reliable diagnosis and treatment of DSRCT patients is required^94^. In addition, access to these technologies is usually restricted to developed countries, which may further impede accurate and timely diagnosis and treatment of DSRCT in economically disadvantaged regions. Thus, there is an urgent need for the development of resources to better address the challenges associated with both the diagnosis and management of DSRCT. While performing comprehensive IHC panels may be laborious and costly, our findings reveal the IHC conditions for optimal CACNA2D2 staining quality and confirmed that single CACNA2D2 staining offers a sensitive, simple, rapid, and economical approach for accurate and reliable diagnosis of DSRCT. In addition, this high, specific and homogenous CACNA2D2 membranous expression in DSRCT in combination with the highly-specific antibody described here, makes CACNA2D2 an ideal candidate for targeted therapeutic approaches including drug delivery using antibody-drug-conjugates or CAR-T cell therapy^95–97^.

In conclusion, we here present a broad toolbox for DSRCT researchers and clinicians: first, a comprehensive multi-omic datasets including single-cell and proteomic data of in vivo generated KD models for EWSR1::WT1. Second, the establishment of a systems-biology-based pipeline combining in silico, in vitro, and in situ analyses to identify strong candidates amenable to validation as diagnostic markers. Third, we computed robust DSRCT-specific gene signatures (both for CACNA2D2 and EWSR1::WT1) that can be used to identify DSRCT cells at bulk and single-cell levels. Lastly, our data collectively support the notion that swift and accurate diagnosis of DSRCT can be facilitated by immunohistochemical detection of the EWSR1::WT1 direct target CACNA2D2, from both FFPE or fresh-frozen samples, which can accurately detect even (micro)metastatic lesions which may still present after chemotherapy and complete cytoreductive treatment in DSRCT patients. As such, we propose to employ CACNA2D2 as a novel single biomarker for DSRCT diagnosis, emphasizing the importance of validating its diagnostic utility in prospective, multi-centered studies.

## ACKNOWLEDGEMENTS

We would like to thank Nadine Gmelin, Stefanie Kutschmann and Felina Zahnow for their expert technical assistance, and Claudia Schmidt from the Light Microscopy Facility (German Cancer Research Center (DKFZ), Heidelberg, Germany) for her meticulous work in conducting immunohistochemical stainings. We thank the Microarray Core Facility (German Cancer Research Center (DKFZ)) for providing the Gene Expression Arrays and related services. We thank Katharina Bauer, Denise Keitel and Jan-Philipp Mallm from the Single-cell Open Lab (German Cancer Research Center (DKFZ)) for expert support in the preparation of single-cell libraries. We thank the Flow Cytometry Facility team (German Cancer Research Centre (DKFZ)) for their support with cell sorting. We thank Dr. Marc Ladanyi for sharing the SK-DSRCT2 cell line.

## AUTHOR CONTRIBUTIONS

F.H.G, F.C.A., and T.G.P.G. conceived the study. F.H.G and F.C.A. wrote the paper, and drafted all figures and tables. F.H.G. carried out all in vitro and in vivo experiments and performed all bioinformatic and statistical analyses. F.H.G., A.R. and T.G.P.G. performed immunohistochemical evaluation and scoring of tumor samples and TMAs. F.C.A, R.I., and A.B. performed and/or coordinated in vivo experiments. O.D. provided microarray expression data. S.K.G. performed in vitro experiments on BER cell lines. T.F. performed single-cell bioinformatic analyses. K.A. and A.O. performed MassSpec and analyzed MassSpec data. A.R., J.L., E.V., L.R.P., M.S. and S.O. contributed to experimental procedures. W.H. and B.F.B.M provided clinical and/or histological guidance. E.A., S.B., S.P.V., M.E., M.S.S., D.B., C.N, D.H., Y.V.J., A.J., S.S. and T.K. provided clinical samples. P.J.G., T.G.P.G., and J.K. provided laboratory infrastructure. F.C.A. and T.G.P.G. supervised the study and data analysis. All authors read and approved the final manuscript.

## SUPPLEMENARY FIGURE LEGENDS

**Supplementary Fig. 1 | CACNA2D2 is significantly overexpressed in DSRCT compared to cancer entities of differential diagnostic relevance**

**A.** Western blot using antibodies against CACNA2D2, WT1, and GAPDH (loading control) in DSRCT tumor samples derived from orthotopic xenografts of JN-DSRCT-1 and SK-DSRCT2 cell line expressing a DOX-inducible shRNA mediated knock down system of EWSR1::WT1. Mice were treated either with DOX or sucrose (control) for 96 h.

**Supplementary Fig. 2 | *CACNA2D2* is directly regulated by a EWSR1::WT1-bound super-enhancer**

**A.** Bar plot showing relative mRNA expression levels of *CACNA2D2* as quantified by qRT-PCR in SK-DSRCT2-endo-WT1-HaloTag cells carrying a knock in of HaloTag-HiBiT tag to the endogenous locus of *EWSR1::WT1*. Horizontal bars represent mean expression levels and whiskers SEM. n≥4 biologically independent experiments. Unpaired two-sided Mann-Whitney test**. B**. Diagram depicting a representation of the different EWSR1::WT1 transcript variants expressed in JN-DSRCT-1, SK-DSRCT2 and BER DSRCT cell lines. The fusion breakpoint encompasses different exons of the *EWSR1* gene. The *WT1* gene either lacks (-KTS) or contains (+KTS) the KTS motif at the junction between exon 9 and 10. **C**. UCSC genome browser showing the epigenetic profile of the *CACNA2D2* locus (chr3:50,506,517-50,548,707) in MeT-5A human mesothelial cells either expressing different isoforms of wild-type WT1 or of EWSR1::WT1. ChIP-seq data derived from GSE212977 are depicted for EWSR1::WT1 (blue), H3K27ac (orange), and WT1 (green). CACNA2D2 promoter and enhancer regions are highlighted with dotted squares. **D**. x-y scatterplot showing H3K27ac signal density in reads per million mapped reads per base at active enhancer sites in MeT-5A cells expressing EWSR1::WT1^−KTS+KTS^ ranked to their normalized intensity from low to high. Horizontal dashed red line indicates cut-off value of 10,645.1 for identification of super-enhancers (n=1,594 as indicated by vertical dashed red line) using ROSE algorithm. **E**. Volcano plots depicting results of pairwise differential gene expression analysis of RNA-seq gene expression profiles from MeT-5A and LP-9 human mesothelial cell lines expressing different isoforms of EWSR1::WT1. Blue dots represent genes with |log2FC|>1.0 and a *Padj*<0.05 (Benjamini-Hochberg corrected).

**Supplementary Fig. 3 | CACNA2D2 is a key component of the EWSR1::WT1 oncogenic signature**

**A.** x-y-scatter plots illustrating correlation of ssGSEA enrichment scores between (sc)EWSR1::WT1 and (sc)CACNA2D2 signatures of orthotopically xenografted metastatic single DSRCT cells. Displayed numbers and colors in the correlation matrix represent Pearson correlation values for each signature enrichment. **B**. Venn diagram showing the number of shared genes between scEWSR1::WT1 and EWSR1::WT1 signatures highlighted in red. **C**. Venn diagram showing the number of shared genes between scEWSR1::WT1, EWSR1::WT1, scCACNA2D2 and CACNA2D2 signatures highlighted in blue. **D**. x-y-scatter plots illustrating correlation of ssGSEA enrichment scores between exclusive (sc)EWSR1::WT1 and (sc)CACNA2D2 signatures of orthotopically xenografted primary and metastatic single DSRCT cells. Displayed numbers and colors in the correlation matrix represent Pearson correlation values for each signature enrichment. **E**. UMAP plot of single-cell RNA-seq analysis of merged and integrated data from eleven samples, comprising tumor-derived and normal cells from four DSRCT patients (GSE263523), using the CACNA2D2 signature. Color gradient indicates the ssGSEA score for CACNA2D2 signature enrichment. CAFs: Cancer-associated fibroblasts. **F**. Violin plot depicting CACNA2D2 signature scores for DSRCT and other normal cell types included in the patient samples. Horizontal bar represents median. Cancer-associated fibroblasts (CAFs).

**Supplementary Fig. 4 | High CACNA2D2 protein expression is a diagnostically-powerful hallmark of DSRCT**

**A.** Boxplot showing mean methylation of *CACNA2D2*-associated CpG sites across 14 cancer entities, including DSRCT. Whiskers indicate minimum and maximum value and horizontal line the median. Number of analyzed samples is given in parentheses. Unpaired two-sided Mann-Whitney test**. B.** UMAP plots of 14 sarcoma types including DSRCT according to each indicated gene’s associated CpG sites. Number of analyzed CpG sites is disclosed in the right bottom corner of each plot. **C.** Diagram depicting the curated (yellow background) or newly generated (blue background) datasets in this study, along with the developed data analysis pipeline. Color of text boxes depicts the origin of the resource (in situ, in vivo, in vitro). Abbreviations: JN-DSRCT-1 or SK-DSRCT-2 cell line models with shRNA mediated KD of EWSR1::WT1 (JN-DSRCT-1^shWT1^, SK-DSRCT-2 ^shWT1^); MeT-5A or LP-9 mesothelial cell line models with ectopic overexpression of either EWSR1::WT1^−KTS^, EWSR1::WT1^+KTS^, or EWSR1::WT1^+/−KTS^ (MeT-5A^EWSR1::WT1^, LP-9^EWSR1::WT1^); BER cell line model transfected with siRNAs targeting *EWSR1::WT1* (BER^siWT1^).

## SUPPLEMENARY TABLES

**Suppl. Table 1 | Oligo nucleotide sequences used in this study**

**Suppl. Table 2 | GEO series and accession codes for ChIP-seq data showing in Figure 2 and Supplementary Figure 2**

**Suppl. Table 3 | Accession codes for ChIP-seq datasets used in ROSE super-enhancer analysis**

**Suppl. Table 4 | Single-cell RNA-seq data accession codes**

**Suppl. Table 5 | Accession codes for all microarray datasets from normal and cancer tissue samples**

**Suppl. Table 6 | DEP analysis of JN-DSRCT-1 and SK-DSRCT2 upon KD of EWSR1::WT1**

**Suppl. Table 7 | ROSE super-enhancer analysis results for JN-DSRCT-1 and MeT-5A cell lines**

**Suppl. Table 8 | *CACNA2D2* gene set**

**Suppl. Table 9 | CACNA2D2 signature, EWSR1::WT1 signature, scCACNA2D2 signature and scEWSR1::WT1 signature**

**Suppl. Table 10 | Solid tumor tissue collection overview**

## REFERENCES

1. Cidre-Aranaz, F. et al. Small round cell sarcomas. Nat. Rev. Dis. Primer 8, 1–22 (2022).

2. de Alava, E. & Marcilla, D. Birth and evolution of the desmoplastic small round-cell tumor. Semin. Diagn. Pathol. 33, 254–261 (2016).

3. WHO Classification of Tumours Editorial Board. Paediatric Tumours. (International agency for research on cancer, Lyon (France), 2022).

4. Ladanyi, M. & Gerald, W. Fusion of the EWS and WT1 genes in the desmoplastic small round cell tumor. Cancer Res. 54, 2837–2840 (1994).

5. Gedminas, J. M. et al. Desmoplastic small round cell tumor is dependent on the EWS-WT1 transcription factor. Oncogenesis 9, 1–8 (2020).

6. Gerald, W. L. et al. Intra-abdominal desmoplastic small round-cell tumor. Report of 19 cases of a distinctive type of high-grade polyphenotypic malignancy affecting young individuals. Am. J. Surg. Pathol. 15, 499–513 (1991).

7. Magrath, J. W. et al. Comprehensive Transcriptomic Analysis of EWSR1::WT1 Targets Identifies CDK4/6 Inhibitors as an Effective Therapy for Desmoplastic Small Round Cell Tumors. Cancer Res. 84, 1426–1442 (2024).

8. Ogura, K. et al. Therapeutic Potential of NTRK3 Inhibition in Desmoplastic Small Round Cell Tumor. Clin. Cancer Res. Off. J. Am. Assoc. Cancer Res. 27, 1184–1194 (2021).

9. Palmer, R. E. et al. Induction of BAIAP3 by the EWS-WT1 chimeric fusion implicates regulated exocytosis in tumorigenesis. Cancer Cell 2, 497–505 (2002).

10. Magrath, J. W. et al. Transcriptomic analysis identifies B-lymphocyte kinase as a therapeutic target for desmoplastic small round cell tumor cancer stem cell-like cells. Oncogenesis 13, 1–15 (2024).

11. Li, H. et al. Adenosine transporter ENT4 is a direct target of EWS/WT1 translocation product and is highly expressed in desmoplastic small round cell tumor. PloS One 3, e2353 (2008).

12. Saito, T. EWS-WT1 Chimeric Protein in Desmoplastic Small Round Cell Tumor is a Potent Transactivator of FGFR4. J. Cancer Sci. Ther. 04, (2012).

13. Negri, T. et al. New transcriptional-based insights into the pathogenesis of desmoplastic small round cell tumors (DSRCTs). Oncotarget 8, 32492–32504 (2017).

14. Hingorani, P. et al. Transcriptome analysis of desmoplastic small round cell tumors identifies actionable therapeutic targets: a report from the Children’s Oncology Group. Sci. Rep. 10, 12318 (2020).

15. Henon, C. et al. Single-cell multiomics profiling reveals heterogeneous transcriptional programs and microenvironment in DSRCTs. Cell Rep. Med. 5, 101582 (2024).

16. Slotkin, E. K. et al. Comprehensive Molecular Profiling of Desmoplastic Small Round Cell Tumor. Mol. Cancer Res. MCR 19, 1146–1155 (2021).

17. Lamhamedi-Cherradi, S.-E. et al. The androgen receptor is a therapeutic target in desmoplastic small round cell sarcoma. Nat. Commun. 13, 3057 (2022).

18. Bleijs, M., et al. EWSR1-WT1 Target Genes and Therapeutic Options Identified in a Novel DSRCT In Vitro Model. Cancers 13, 6072 (2021).

19. Gerald, W. L. & Rosai, J. Case 2. Desmoplastic small cell tumor with divergent differentiation. Pediatr. Pathol. 9, 177–183 (1989).

20. Markides, C. S. A. et al. Desmoplastic small round cell tumor (DSRCT) xenografts and tissue culture lines: Establishment and initial characterization. Oncol. Lett. 5, 1453–1456 (2013).

21. Wiederschain, D. et al. Single-vector inducible lentiviral RNAi system for oncology target validation. Cell Cycle Georget. Tex 8, 498–504 (2009).

22. Musa, J. et al. Cooperation of cancer drivers with regulatory germline variants shapes clinical outcomes. Nat. Commun. 10, 4128 (2019).

23. Los, G. V. et al. HaloTag: A Novel Protein Labeling Technology for Cell Imaging and Protein Analysis. ACS Chem. Biol. 3, 373–382 (2008).

24. Schwinn, M. K. et al. CRISPR-Mediated Tagging of Endogenous Proteins with a Luminescent Peptide. ACS Chem. Biol. 13, 467–474 (2018).

25. Cidre-Aranaz, F. et al. Integrative gene network and functional analyses identify a prognostically relevant key regulator of metastasis in Ewing sarcoma. Mol. Cancer 21, 1 (2022).

26. Heinz, S. et al. Simple combinations of lineage-determining transcription factors prime cis-regulatory elements required for macrophage and B cell identities. Mol. Cell 38, 576–589 (2010).

27. Church, D. M. et al. Modernizing Reference Genome Assemblies. PLOS Biol. 9, e1001091 (2011).

28. Grant, C. E., Bailey, T. L. & Noble, W. S. FIMO: scanning for occurrences of a given motif. Bioinforma. Oxf. Engl. 27, 1017–1018 (2011).

29. Sayers, E. W. et al. Database resources of the national center for biotechnology information. Nucleic Acids Res. 50, D20–D26 (2022).

30. Li, H. Aligning sequence reads, clone sequences and assembly contigs with BWA-MEM. Preprint at 10.48550/arXiv.1303.3997 (2013).

31. Danecek, P. et al. Twelve years of SAMtools and BCFtools. GigaScience 10, giab008 (2021).

32. Zhang, Y. et al. Model-based Analysis of ChIP-Seq (MACS). Genome Biol. 9, R137 (2008).

33. Whyte, W. A. et al. Master Transcription Factors and Mediator Establish Super-Enhancers at Key Cell Identity Genes. Cell 153, 307–319 (2013).

34. Lovén, J. et al. Selective Inhibition of Tumor Oncogenes by Disruption of Super-Enhancers. Cell 153, 320–334 (2013).

35. Truong, D. D. et al. Dissociation protocols used for sarcoma tissues bias the transcriptome observed in single-cell and single-nucleus RNA sequencing. BMC Cancer 23, 488 (2023).

36. Picelli, S. et al. Full-length RNA-seq from single cells using Smart-seq2. Nat. Protoc. 9, 171–181 (2014).

37. Hao, Y. et al. Dictionary learning for integrative, multimodal and scalable single-cell analysis. Nat. Biotechnol. 42, 293–304 (2024).

38. Hao, Y. et al. Integrated analysis of multimodal single-cell data. Cell 184, 3573–3587.e29 (2021).

39. Stuart, T. et al. Comprehensive Integration of Single-Cell Data. Cell 177, 1888–1902.e21 (2019).

40. Butler, A., Hoffman, P., Smibert, P., Papalexi, E. & Satija, R. Integrating single-cell transcriptomic data across different conditions, technologies, and species. Nat. Biotechnol. 36, 411–420 (2018).

41. Satija, R., Farrell, J. A., Gennert, D., Schier, A. F. & Regev, A. Spatial reconstruction of single-cell gene expression data. Nat. Biotechnol. 33, 495–502 (2015).

42. Amezquita, R. A. et al. Orchestrating single-cell analysis with Bioconductor. Nat. Methods 17, 137–145 (2020).

43. Hippen, A. A. et al. miQC: An adaptive probabilistic framework for quality control of single-cell RNA-sequencing data. 2021.03.03.433798 Preprint at 10.1101/2021.03.03.433798 (2021).

44. McCarthy, D. J., Campbell, K. R., Lun, A. T. L. & Wills, Q. F. Scater: pre-processing, quality control, normalization and visualization of single-cell RNA-seq data in R. Bioinformatics 33, 1179–1186 (2017).

45. Eisenberg, E. & Levanon, E. Y. Human housekeeping genes, revisited. Trends Genet. TIG 29, 569–574 (2013).

46. Adam, M., Potter, A. S. & Potter, S. S. Psychrophilic proteases dramatically reduce single-cell RNA-seq artifacts: a molecular atlas of kidney development. Dev. Camb. Engl. 144, 3625–3632 (2017).

47. van den Brink, S. C., et al. Single-cell sequencing reveals dissociation-induced gene expression in tissue subpopulations. Nat. Methods 14, 935–936 (2017).

48. Denisenko, E. et al. Systematic assessment of tissue dissociation and storage biases in single-cell and single-nucleus RNA-seq workflows. Genome Biol. 21, 130 (2020).

49. Fan, C. et al. irGSEA: the integration of single-cell rank-based gene set enrichment analysis. Brief. Bioinform. 25, bbae243 (2024).

50. Baldauf, M. C. et al. Robust diagnosis of Ewing sarcoma by immunohistochemical detection of super-enhancer-driven EWSR1-ETS targets. Oncotarget 9, 1587–1601 (2017).

51. Surdez, D. et al. Targeting the EWSR1-FLI1 Oncogene-Induced Protein Kinase PKC-β Abolishes Ewing Sarcoma Growth. Cancer Res. 72, 4494–4503 (2012).

52. Gautier, L., Cope, L., Bolstad, B. M. & Irizarry, R. A. affy--analysis of Affymetrix GeneChip data at the probe level. Bioinforma. Oxf. Engl. 20, 307–315 (2004).

53. Irizarry, R. A. et al. Exploration, normalization, and summaries of high density oligonucleotide array probe level data. Biostat. Oxf. Engl. 4, 249–264 (2003).

54. Dai, M. et al. Evolving gene/transcript definitions significantly alter the interpretation of GeneChip data. Nucleic Acids Res. 33, e175 (2005).

55. Ritchie, M. E. et al. limma powers differential expression analyses for RNA-sequencing and microarray studies. Nucleic Acids Res. 43, e47 (2015).

56. Love, M. I., Huber, W. & Anders, S. Moderated estimation of fold change and dispersion for RNA-seq data with DESeq2. Genome Biol. 15, 550 (2014).

57. Korotkevich, G. et al. Fast gene set enrichment analysis. bioRxiv 060012 (2021) doi:10.1101/060012.

58. Subramanian, A. et al. Gene set enrichment analysis: A knowledge-based approach for interpreting genome-wide expression profiles. Proc. Natl. Acad. Sci. 102, 15545–15550 (2005).

59. Liberzon, A. et al. Molecular signatures database (MSigDB) 3.0. Bioinformatics 27, 1739– 1740 (2011).

60. Liberzon, A. et al. The Molecular Signatures Database (MSigDB) hallmark gene set collection. Cell Syst. 1, 417–425 (2015).

61. Milacic, M. et al. The Reactome Pathway Knowledgebase 2024. Nucleic Acids Res. 52, D672–D678 (2024).

62. Gu, Z., Eils, R. & Schlesner, M. Complex heatmaps reveal patterns and correlations in multidimensional genomic data. Bioinformatics 32, 2847–2849 (2016).

63. Ewing Sarcoma: Methods and Protocols. vol. 2226 (Springer US, New York, NY, 2021).

64. Baldauf, M. C. et al. Systematic identification of cancer-specific MHC-binding peptides with RAVEN. Oncoimmunology 7, e1481558 (2018).

65. Marchetto, A. et al. Oncogenic hijacking of a developmental transcription factor evokes vulnerability toward oxidative stress in Ewing sarcoma. Nat. Commun. 11, 2423 (2020).

66. Dallmayer, M. et al. Targeting the CALCB/RAMP1 axis inhibits growth of Ewing sarcoma. Cell Death Dis. 10, 116 (2019).

67. Ohmura, S. et al. Translational evidence for RRM2 as a prognostic biomarker and therapeutic target in Ewing sarcoma. Mol. Cancer 20, 97 (2021).

68. Müller, T. et al. Automated sample preparation with SP3 for low-input clinical proteomics. Mol. Syst. Biol. 16, e9111 (2020).

69. Müller, T., Cremonini, M. A., Kliewer, G. & Krijgsveld, J. Automated Sample Preparation for Mass Spectrometry-Based Clinical Proteomics. in Mass Spectrometry-Based Proteomics (ed. Gevaert, K.) 181–211 (Springer US, New York, NY, 2023). doi:10.1007/978-1-0716-3457-8_11.

70. Demichev, V., Messner, C. B., Vernardis, S. I., Lilley, K. S. & Ralser, M. DIA-NN: neural networks and interference correction enable deep proteome coverage in high throughput. Nat. Methods 17, 41–44 (2020).

71. Koelsche, C. et al. Sarcoma classification by DNA methylation profiling. Nat. Commun. 12, 498 (2021).

72. McInnes, L., Healy, J., Saul, N. & Großberger, L. UMAP: Uniform Manifold Approximation and Projection. J. Open Source Softw. 3, 861 (2018).

73. McInnes, L., Healy, J. & Melville, J. UMAP: Uniform Manifold Approximation and Projection for Dimension Reduction. Preprint at 10.48550/arXiv.1802.03426 (2020).

74. Wickham, H. Data Analysis. in ggplot2 189–201 (Springer International Publishing, Cham, 2016). doi:10.1007/978-3-319-24277-4_9.

75. Thway, K. et al. Desmoplastic Small Round Cell Tumor: Pathology, Genetics, and Potential Therapeutic Strategies. Int. J. Surg. Pathol. 24, 672–684 (2016).

76. Grünewald, T. G. P. et al. Ewing sarcoma. Nat. Rev. Dis. Primer 4, 5 (2018).

77. Nishio, J. et al. Establishment and Characterization of a Novel Human Desmoplastic Small Round Cell Tumor Cell Line, JN-DSRCT-1. Lab. Invest. 82, 1175–1182 (2002).

78. Smith, R. S. et al. Novel patient-derived models of desmoplastic small round cell tumor confirm a targetable dependency on ERBB signaling. Dis. Model. Mech. 15, dmm047621 (2022).

79. Gryder, B. E. et al. PAX3-FOXO1 Establishes Myogenic Super Enhancers and Confers BET Bromodomain Vulnerability. Cancer Discov. 7, 884–899 (2017).

80. Riggi, N. et al. EWS-FLI1 utilizes divergent chromatin remodeling mechanisms to directly activate or repress enhancer elements in Ewing sarcoma. Cancer Cell 26, 668–681 (2014).

81. Hnisz, D. et al. Super-Enhancers in the Control of Cell Identity and Disease. Cell 155, 934– 947 (2013).

82. Antonescu, C. R., Gerald, W. L., Magid, M. S. & Ladanyi, M. Molecular variants of the EWS-WT1 gene fusion in desmoplastic small round cell tumor. Diagn. Mol. Pathol. Am. J. Surg. Pathol. Part B 7, 24–28 (1998).

83. Hartono, A. B. et al. Salt-Inducible Kinase 1 is a potential therapeutic target in Desmoplastic Small Round Cell Tumor. Oncogenesis 11, 1–10 (2022).

84. Hayes-Jordan, A., LaQuaglia, M. P. & Modak, S. Management of desmoplastic small round cell tumor. Semin. Pediatr. Surg. 25, 299–304 (2016).

85. Remmele, W. & Stegner, H. E. [Recommendation for uniform definition of an immunoreactive score (IRS) for immunohistochemical estrogen receptor detection (ER-ICA) in breast cancer tissue]. Pathol. 8, 138–140 (1987).

86. Li, J. et al. Therapeutic targeting of the PLK1-PRC1-axis triggers cell death in genomically silent childhood cancer. Nat. Commun. 12, 5356 (2021).

87. Sannino, G. et al. Gene expression and immunohistochemical analyses identify SOX2 as major risk factor for overall survival and relapse in Ewing sarcoma patients. EBioMedicine 47, 156–162 (2019).

88. Davis, J. L. & Rudzinski, E. R. Small Round Blue Cell Sarcoma Other Than Ewing Sarcoma: What Should an Oncologist Know? Curr. Treat. Options Oncol. 21, 90 (2020).

89. Trikalinos, N. A., Chrisinger, J. S. A. & Van Tine, B. A. Common Pitfalls in Ewing Sarcoma and Desmoplastic Small Round Cell Tumor Diagnosis Seen in a Study of 115 Cases. Med. Sci. 9, 62 (2021).

90. Thway, K. et al. Desmoplastic Small Round Cell Tumor:Pathology, Genetics, and Potential Therapeutic Strategies. Int. J. Surg. Pathol. 24, 672–684 (2016).

91. Murphy, A. J. et al. A new molecular variant of desmoplastic small round cell tumor: significance of WT1 immunostaining in this entity. Hum. Pathol. 39, 1763–1770 (2008).

92. Nakatsuka, S. et al. Immunohistochemical detection of WT1 protein in a variety of cancer cells. Mod. Pathol. 19, 804–814 (2006).

93. Qi, X. et al. Wilms’ tumor 1 (WT1) expression and prognosis in solid cancer patients: a systematic review and meta-analysis. Sci. Rep. 5, 8924 (2015).

94. Allemani, C. et al. Global surveillance of cancer survival 1995–2009: analysis of individual data for 25 676 887 patients from 279 population-based registries in 67 countries (CONCORD-2). The Lancet 385, 977–1010 (2015).

95. Flynn, P., Suryaprakash, S., Grossman, D., Panier, V. & Wu, J. The antibody–drug conjugate landscape. Nat. Rev. Drug Discov. (2024) doi:10.1038/d41573-024-00064-w.

96. Bagley, S. J. et al. Intrathecal bivalent CAR T cells targeting EGFR and IL13Rα2 in recurrent glioblastoma: phase 1 trial interim results. Nat. Med. 30, 1320–1329 (2024).

97. June, C. H., O’Connor, R. S., Kawalekar, O. U., Ghassemi, S. & Milone, M. C. CAR T cell immunotherapy for human cancer. Science 359, 1361–1365 (2018).

