## Supplementary Figures for "Super-enhancer-driven *CACNA2D2* is an EWSR1::WT1 signature gene encoding a diagnostic marker for desmoplastic small round cell tumor (DSRCT)"

**Supplementary Figure 1** Geyer *et al.*

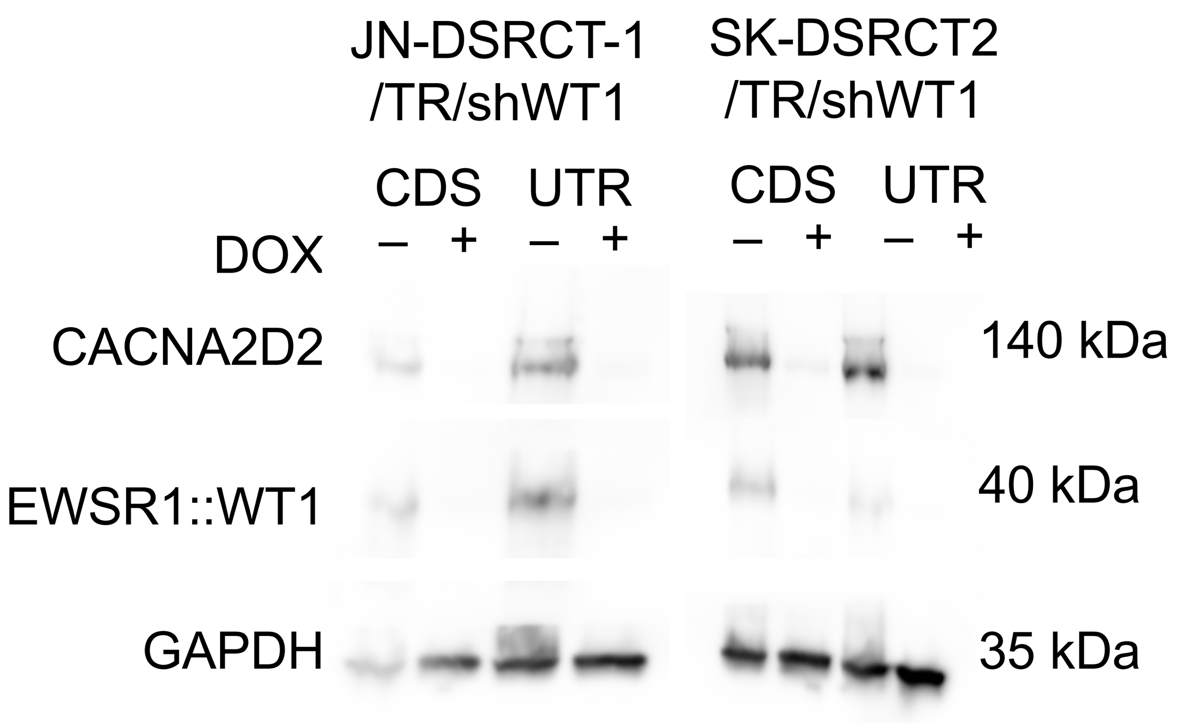

**Supplementary Figure 2** Geyer *et al.*

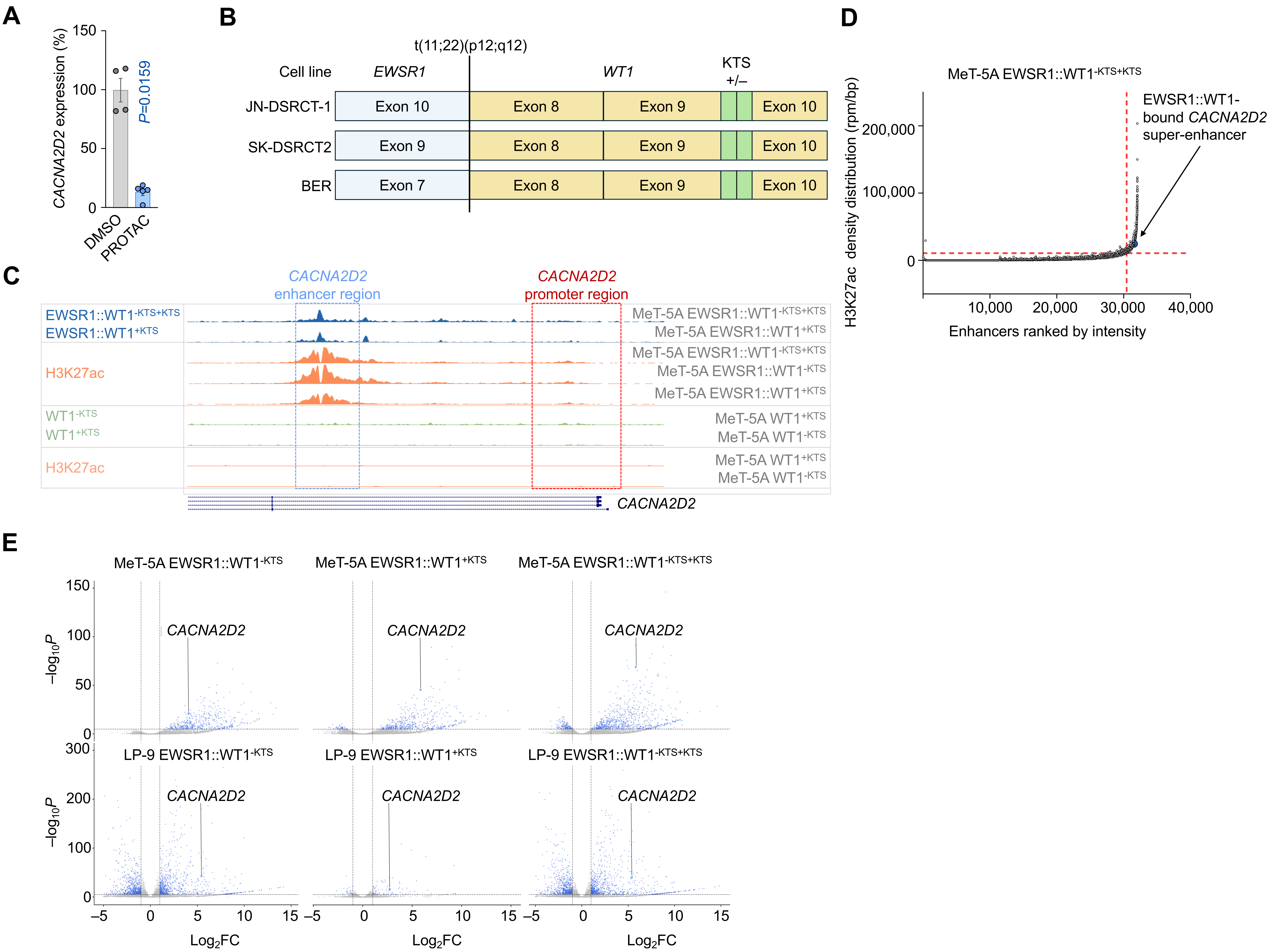

**Supplementary Figure 3** Geyer *et al.*

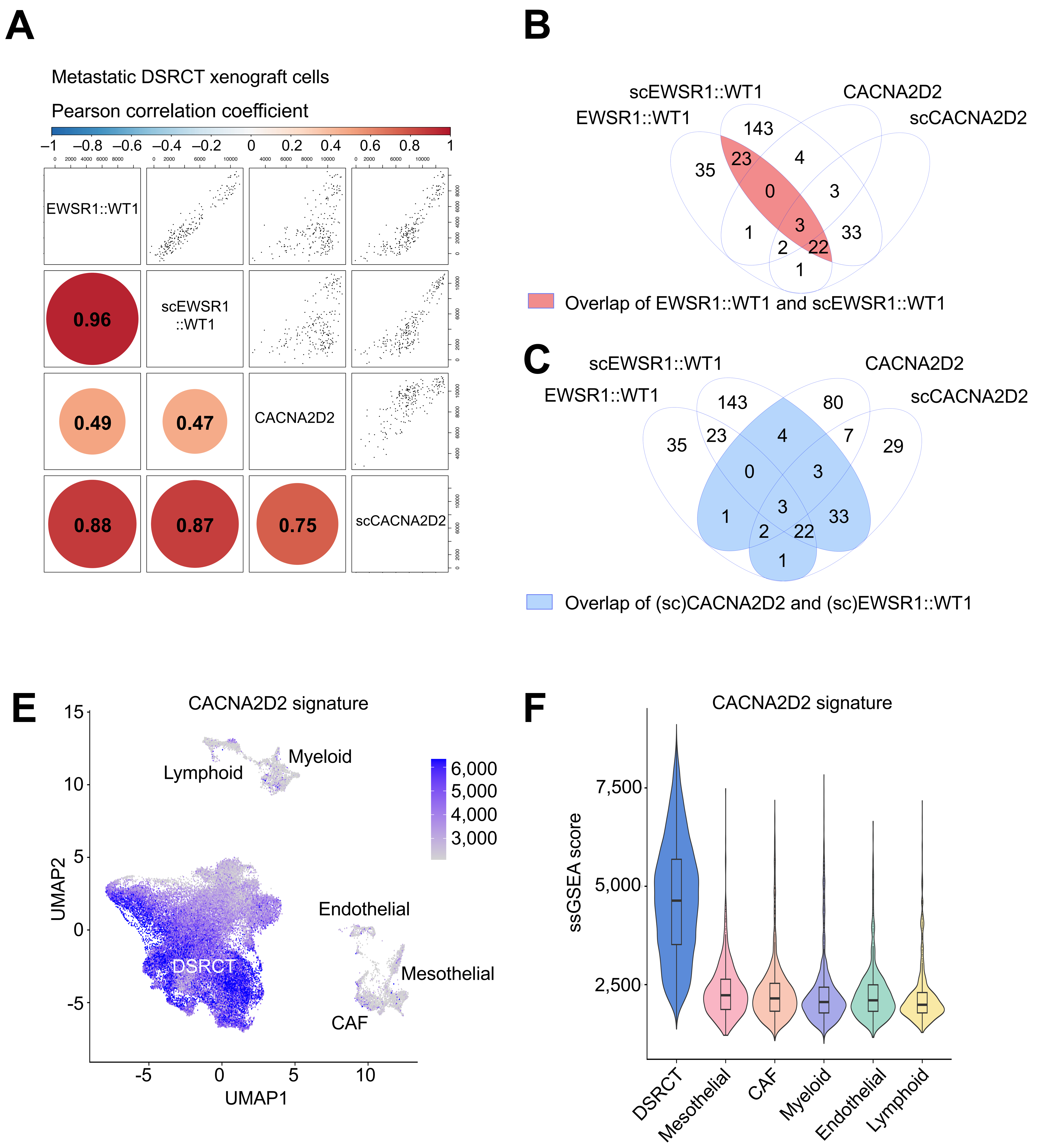

**Supplementary Figure 4** Geyer *et al.*

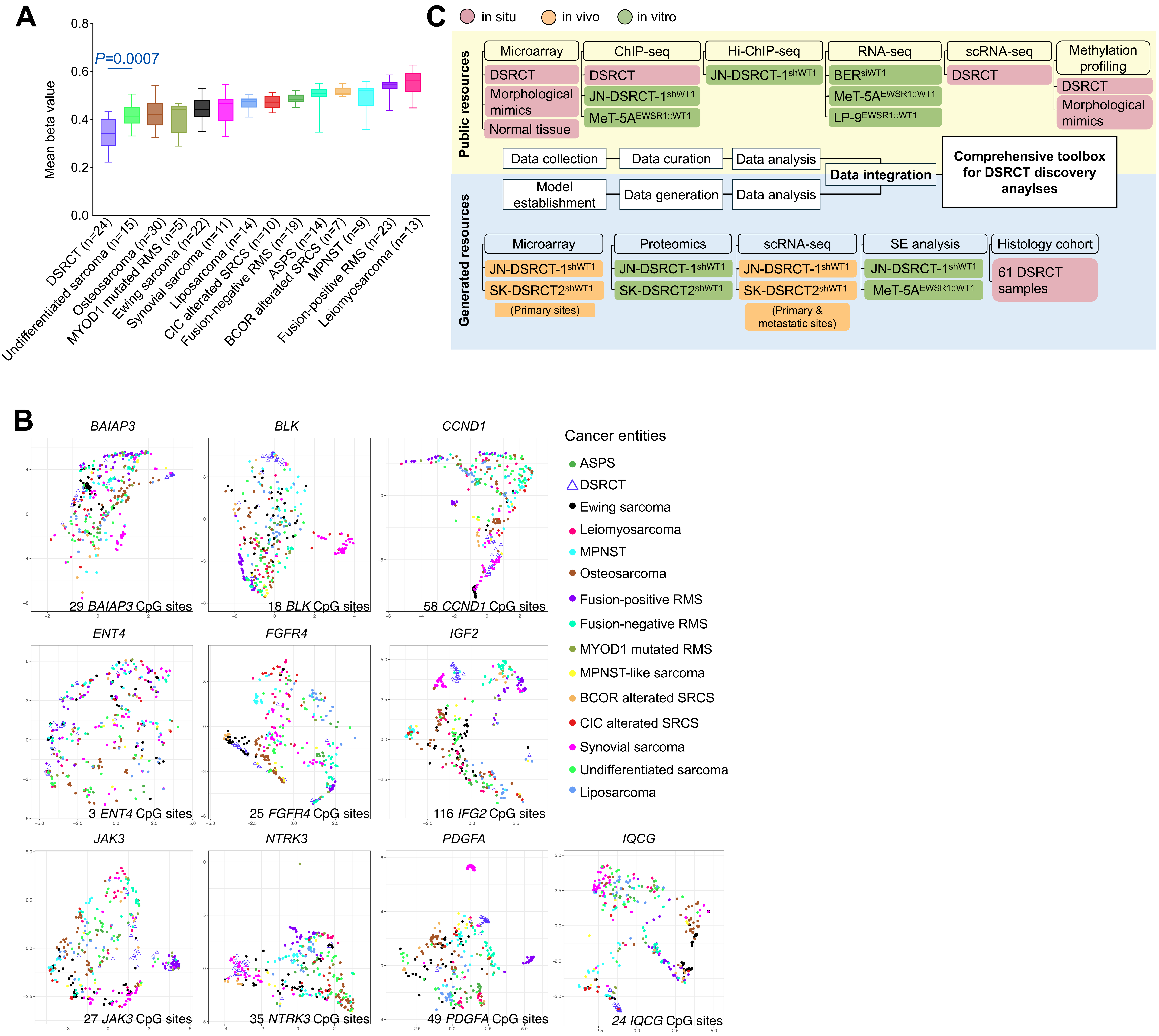
